# Genomic and physiological characterization of *Novosphingobium terrae* sp. nov., an alphaproteobacterium isolated from Cerrado soil containing a megasized chromid

**DOI:** 10.1101/2021.09.13.460193

**Authors:** Aline Belmok, Felipe Marques de Almeida, Rodrigo Theodoro Rocha, Carla Simone Vizzotto, Marcos Rogério Tótola, Marcelo Henrique Soller Ramada, Ricardo Henrique Krüger, Cynthia Maria Kyaw, Georgios J. Pappas

## Abstract

A novel bacterial strain, designated GeG2^T^, was isolated from soils of native Cerrado, a highly biodiverse savanna-like Brazilian biome. 16S rRNA gene sequence analysis of strain GeG2^T^ revealed high sequence identity (100%) to the alphaproteobacterium *Novosphingobium rosa*, however, comparisons with *N. rosa* DSM7285^T^ showed several distinctive features, prompting a full characterization of the new strain in terms of growth, morphology, biochemistry and, ultimately, its genome. GeG2^T^ cells were Gram-stain negative bacilli, facultatively anaerobic, motile, positive for catalase and oxidase activities and for starch hydrolysis. Strain GeG2^T^ presented planktonic-sessile dimorphism and cell aggregates surrounded by extracellular matrix and nanometric spherical structures were observed in liquid cultures, suggesting the production of exopolysaccharides (EPS) and outer membrane vesicles (OMVs). Whole genome assembly revealed four circular replicons: a 4.1 Mb chromosome, a 2.7 Mb extrachromosomal megareplicon and two plasmids (212.7 and 68.6 kb). The megareplicon contains few core genes and plasmid-type replication/maintenance systems, consistent with its classification as a chromid. Genome annotation shows a vast repertoire of carbohydrate active enzymes and genes involved in the degradation of aromatic compounds, highlighting the biotechnological potential of the new isolate obtained from Cerrado soils, especially regarding EPS production and biodegradation of recalcitrant compounds. Chemotaxonomic features, including polar lipid and fatty acid profiles, as well as physiological, molecular and whole genome comparisons showed significant differences between strain GeG2^T^ and a *N. rosa*, clearly indicating that it represents a novel species, for which the name *Novosphingobium terrae* is proposed. The type strain is GeG2^T^ (=CBMAI 2313T =CBAS 753T).

**IMPORTANCE:** *Novosphingobium* is an alphaproteobacterial genus presenting diverse physiological profiles and broad biotechnological applications. However, many aspects regarding the biology of this important bacterial group remain elusive. A novel *Novosphingobium* strain was isolated from soils of Cerrado, an important Brazilian biome. Despite 100% 16S rRNA gene identity with *Novosphingobium rosa*, polyphasic characterizations, including physiological, chemotaxonomic, and whole genome-based analyses revealed significant differences between GeG2^T^ and N. rosa DSM7285^T^, reinforcing resolution limitations in phylogenetic analysis based solely on 16S RNA and highlighting the importance of employing different approaches for the description of bacterial species. Using short and long read sequencing approaches, a high-quality fully resolved genome assembly was generated and one of the largest chromids reported to date was identified. A comprehensive characterization of environmental isolates allows us to better elucidate the diversity and biology of members of this bacterial group with potential biotechnological importance, guiding future bioprospecting efforts and genomic studies.

## INTRODUCTION

*Novosphingobium* is a genus that belongs to the Alphaproteobacteria class, with its members manifesting diverse physiological profiles and displaying the capacity to colonize diverse niches such as soils and rhizospheres (Takeuchi et al. 1995; Kämpfer et al. 2011; Kämpfer et al. 2015), ground water (Lee et al., 2014a), freshwater (Baek et al., 2011; Ngo et al., 2016; Sheu et al., 2016), deep-sea environments (Yuan et al., 2009; Huo et al., 2015), mangrove sediments (Lee et al., 2014b), bioremediation systems (Tiirola et al. 2005), coolant lubricant emulsion (Kämpfer et al. 2018), among many other natural and artificial habitats.

These organisms are Gram-negative non-sporulating small bacilli (1-4 µm x 0.3-1.0 µm), motile or non-motile, aerobic or facultative anaerobic (Glaeser and Kämpfer, 2014). *Novosphingobium* species are known for their production of different exopolysaccharides (EPS), which are commonly used in beverage and food industries (Wu *et al*., 2017), and whose production is affected by environmental and *in vitro* culture conditions. Members of this genus also present the ability to degrade a wide variety of xenobiotics and aromatic compounds, such as polycyclic aromatic hydrocarbons (PAHs), lignin derivatives, heterocyclic compounds, steroid endocrine disruptors, and pesticides (Tiirola et al., 2005; Hegedus et al., 2017; Kampfer et al., 2018; Sheu et al., 2018, Wang et al., 2018), qualifying them as candidates for different bioremediation processes.

At the time of writing, the List of Prokaryotic names with Standing in Nomenclature (LPSN) recognized 52 *Novosphingobium* species (https://lpsn.dsmz.de/genus/novosphingobium). However, despite their metabolic versatility and potential biotechnological and industry applications, little is known about their biology. To date, only 24 complete genomes are available on public databases, reinforcing the lack of information about this important bacterial group.

In this work, we describe the genomic and physiological characterization of a *Novosphingobium* strain (GeG2^T^) isolated from soils of a Brazilian savannah-like biome, known as Cerrado. A comprehensive morphological, chemotaxonomic, molecular, biochemical and genome sequence analysis revealed that, despite sharing identical 16S rRNA sequence with *Novosphingobium rosa,* GeG2^T^ represents a new species, for which the name *Novosphingobium terrae* is proposed. Its environmental adaptation repertoire can be assessed by the existence of a varied set of genes related to polysaccharide catabolism, particularly xylan and hemicellulose, as well as genes conferring the ability to degrade recalcitrant aromatic compounds. Remarkably, in addition to the main chromosome, we could detect and annotate a secondary megabase-sized replicon, most likely a chromid (Harrison et al., 2010), one of the largest reported to date.

## MATERIAL AND METHODS

### Bacteria Isolation and Cultivation

The microorganism was isolated from soils sampled at a native area of Cerrado, a savannah-like Brazilian biome, located at the Reserva Ecológica do IBGE, Brasília, Brazil (15°55’ S, 47°51’ W), in January 2014. The isolate, denominated strain GeG2^T^, was initially enriched in solid media consisting of soil extracts (5% - w/v) and agar (1,5% - w/v), supplemented with ampicillin (150 μg/mL), streptomycin (50 μg/mL), chloramphenicol (20 μg/mL) and itraconazole (0.25 mg/mL). Plates were incubated at 28 °C and cultures were transferred into fresh media approximately once a month. For strain GeG2^T^ isolation, colonies were transferred to minimal medium (MM, per liter: 0.1 g KH2PO4; 0.2 g (NH4)2SO4; 0.1 g MgSO4.7H2O; 0.02 g CaCl2.2H2O; 0.2 g NaCl; 0.1 g yeast extract; 0.05% glucose; 15 g bacteriological agar, pH 5.5), using the standard streak plate technique to obtain pure cultures. Plates were incubated at 28 °C for 48 hours and strain GeG2^T^ was stored at -80 °C in sterile MM containing 20% (v/v) glycerol.

### 16S rRNA gene phylogenetic analysis

The 16S rRNA gene of strain GeG2^T^ was amplified by polymerase chain reactions (PCR) using universal bacterial primers 27F (5‘AGAGTTTGATCCTGGCTCAG3’)/1492R (5‘GGTTACCTTGTTACGACTT3’) (Lane, 1991). PCR reactions were performed in a Bio Rad PTC- 100® (Peltier Thermal Cycler) employing the following cycle conditions: 5 min at 95 °C, followed by 30 cycles of 1 min at 95 °C, 1 min at 55 °C, 2 min at 72 °C and a final extension of 10 min at 72 °C. PCR products were purified with Wizard^®^ SV Gel and PCR Clean-Up System (Promega) and ligated to the pGEM-T easy^®^ vector (Promega), according to manufacturer’s instructions. Plasmid extraction was performed with phenol-chloroform-isoamyl alcohol at 25:24:1 (v:v:v) and Sanger sequencing was carried out by Macrogen Inc. (Seoul, Korea) using universal primers T7 e SP6. The EzTaxon-e server (available at http://www.ezbiocloud.net) (Yoon et al., 2017) was used to calculate the similarity between 16S rRNA gene sequences of GeG2^T^ and other strains. Phylogenetic trees were constructed by neighbor-joining and maximum-likelihood methods using MEGA 5.0 software (Tamura *et al*., 2011), based on Maximum Composite Likelihood (Tamura *et al*., 2004) and Kimura 2-parameter models (Kimura, 1980), respectively, with reliability confirmed by bootstrap analysis based on 1000 resamplings.

### Physiological and chemotaxonomic characterizations

A *Novosphingobium rosa* strain, DSM7285^T^, was obtained from the Leibniz-Institut Deutsche Sammlung von Mikroorganismen und Zellkulturen (DSMZ) and used as a reference strain for comparative phenotypic analyses against GeG2^T^. Growth of both strains in TSA (Trypticase Soy Agar), LB (Luria-Bertani), NA (Nutrient Agar) and MM media was assessed. Unless mentioned otherwise, for all further characterizations, both bacterial strains were cultured on MM containing glucose (0.05% - w/v) as carbon source, at 28 °C.

Growth of GeG2^T^ and reference strain DSM7285^T^ in different temperatures (4, 15, 20, 28, 33, 37 and 42 °C) and different pH (4.0 to 9.0 in 1.0-unit intervals, buffered with 50 mM MES - pH 4.0 to 6.0; MOPS - pH 7.0 or Tris - pH 8.0 and 9.0) was evaluated in agar plates incubated for one week. Salinity requirement and tolerance were evaluated on solid MM supplemented with 0, 0.1, 0.3, 0.5, 0.8, 1.0, 1.5, 2.0 and 3.0% (w/v) of NaCl. Hydrolysis of starch was analyzed in solid MM supplemented with soluble starch (0.5% - w/v) and revealed with iodine vapor after 7, 14 and 24 days (Tindall et al., 2007). Catalase activity was determined by assessing bubble production by cells in 3% (v/v) H2O2. Growth under anaerobic conditions was tested in agar medium supplemented with cysteine hydrochloride (0.05% - w/v), sodium sulfide (0.05% - w/v), and potassium nitrate (0.1% - w/v). Anaerobic glass jars were prepared under a nitrogen atmosphere, sealed with rubber stoppers and aluminum seals, and incubated at 28 °C for 35 days. Jars maintained under aerobic conditions were used as controls. Nitrate reduction, indole production, urease and gelatinase tests, assimilation and oxidation of various carbon compounds and enzyme activities were carried out by the Identification Service of DSMZ Leibiniz Institute, using the API 20NE and API ZYM kits (bioMérieux), according to manufacturer’s instructions.

Antibiotic susceptibility profiles of strain GeG2^T^ were evaluated by the agar-diffusion method using antibiotic-impregnated discs (Bauer et al., 1966, Sha et al., 2017), with bacterial suspensions spread over MM plates, incubated at 28 °C for 48 hours. The tested antibiotics were nalidixic acid (30 μg), amikacin (30 μg), amoxicillin + clavulanic acid (20 + 10 μg), ampicillin (10 μg), cephalothin (30 μg), cefepime (30 μg), cefoxitin (30 μg), ceftazidime (30 μg), cefuroxime (30 μg), ciprofloxacin (5 μg), clindamycin (2 μg), chloramphenicol (30 μg), erythromycin (15 μg), gentamicin (10 μg), levofloxacin (5 μg), meropenem (10 μg), nitrofurantoin (300 μg), norfloxacin (10 μg), oxacillin (1 μg), penicillin (6 μg), rifampicin (5 μg), tetracycline (30 μg), trimethoprim + sulfamethoxazole (1.25 + 23.75 μg), and vancomycin (30 μg).

Fatty acid compositions of strains GeG2^T^ and DSM7285^T^ grown in solid MM for 72 hours were determined following the standard protocol of Sherlock Microbial Identification System (version 6.2) (Sasser, 1990). Briefly, approximately 40 mg of bacterial biomass harvested from third quadrants were submitted to saponification in 1 mL methanol/sodium hydroxide solution (150 mL deionized water, 150 mL methanol, 45 g sodium hydroxide), followed by methylation in 2 mL of 6 mol/L methane in HCl and extraction with 1.25 mL hexane:tertbutyl ether (1:1). The fatty acid profiles were analyzed by gas chromatography (Agilent 7890A) using the RTSBA6 method/library. Analysis of strain GeG2^T^ polar lipids and respiratory quinones were performed by the Identification Services of DSMZ Leibiniz Institute, following standard protocols (Tindall, 1990a; Tindall, 1990b; Tindall et al., 2007).

### Protein profiles analyses by MALDI-TOF

Protein profiles of strain GeG2^T^ and the reference strain DSM7285^T^ were determined by MALDI-TOF mass spectrometry. Proteins were extracted using the protocol described by Agustini et al., 2014 and then spotted (1 μL of protein extraction) in technical sextuplicates on a 96-well target steel plate, covered by 1 μL of 10 mg/mL α-cyano-4-hydroxycinnamic acid (HCCA) matrix (50% (v/v) acetonitrile, 0.3% (v/v) trifluoroacetic acid) and let it dry at room temperature. Spectra were obtained in an Autoflex Speed II MALDI-TOF/TOF (Bruker Daltonics) in positive linear mode, 2,000–20,000 m/z range, acquiring 2,000 successful shots per spot using FlexControl 3.0 software. In total, 10,000 laser shots were accumulated for each spectrum. Spectra were further analyzed in MALDI Biotyper 3.0 software (Ramasamy et al., 2014) that compares each sample mass spectrum to reference mass spectra in the Biotyper database and calculates a score value between 0 and 3 reflecting their similarity. GeG2^T^ and DSM7285^T^ main spectra profiles (MSP) were compared to each other and to Biotyper database. Scores of >2.0 were accepted as reliable for identification at the species level, >1.7 but <2.0 at the genus level and scores <1.7 were considered unreliable, as specified by the manufacturer. All experiments were performed in biological sextuplicate. A dendrogram was constructed using MSP information from each replicate on MALDI Biotyper 3.0 software.

### Microscopy

GeG2^T^ morphological characterizations were performed by light and electronic microscopy analyses. Fresh or Gram-stained cells grown on solid or liquid media were observed by phase-contrast or bright field microscopy, respectively, in a Axio Scope.A1 (Zeiss, Germany) microscope. Scanning electronic microscopy (SEM) was performed in cells grown in liquid media for 72 hours or 14 days and fixed on Karnovsky fixative solution (Souza, 2007). Images were generated with a JEOL JSM- 7001F microscope (JEOL Ltd., Tokyo, Japan). Transmission electron microscopy (TEM) analyses were conducted in the Center of Microscopy at the Universidade Federal de Minas Gerais using bacterial cells grown for 7 days in liquid or solid media and posteriorly cryofixed by high-pressure freezing (HPF) or cells grown for 14 days in liquid media and fixed on Karnovsky fixative solution.

### Genome sequencing and assembly

Genomic DNA was extracted from cultures using a standard phenol-chloroform method. Briefly, cells were harvested from culture plates and resuspended in a lysis buffer containing 25 mM Tris-HCl, 10 mM EDTA, 200 μg/mL proteinase K, 100 μg/mL RNAse A and 0.6% (w/v) SDS. After incubation for 1 hour at 37 °C, an equal volume of phenol:chloroform:isoamyl alcohol mixture (25:24:1) was added. The aqueous phase was then treated with chloroform:isoamyl alcohol (24:1) and DNA was precipitated with 0.3 M NaCl and cold ethanol. After washing with 70% ethanol (v/v), the extracted DNA was dried and re-suspended in ultra-pure H2O.

Genome sequencing was performed at Macrogen Inc., Korea, using a combination of Illumina and PacBio technologies. Illumina libraries were constructed with TrueSeq DNA shotgun PCR-Free (350 bp) kits and Pacbio 20 kb bluepippin systems. Sequencing was performed on Illumina HiSeq 2000 (paired-end sequencing) and PacBio RS (2 SMRT cells) platforms, respectively.

Illumina short reads were preprocessed with FastQC v0.11.5 (Andrews, 2010) and TrimGalore v0.6.0 (https://github.com/FelixKrueger/TrimGalore). PacBio reads were extracted using pbh5tools v0.8.0 (github.com/PacificBiosciences/pbh5tools), using the ngs-preprocess pipeline (github.com/fmalmeida/ngs-preprocess), with default parameters. Additionally, Illumina paired-end reads were merged with PEAR v0.9.8 (Zhang et al., 2014) to increase overall read length and help in the assembly step. Preprocessed Illumina and PacBio reads were assembled together with the reads merged by PEAR using the program Unicycler v0.4.7 (Wick, et al. 2017), in hybrid mode, and assembly statistics was assessed with QUAST v5.0.1 (Gurevich et al. 2013), both part of the MpGAP pipeline (github.com/fmalmeida/MpGAP). Genome completeness was assessed using BUSCO v3.1.0 (Waterhouse et al. 2018) and CheckM v1.0.13 (Parks et al., 2014), using sphingomonadales_odb10 and o_Sphingomonadales (UID3310) reference datasets, respectively. Circular representations for the different genome replicons were generated with DNAPlotter (Carver et al., 2009).

Genome-based taxonomic identification was performed by OGRI (Overall Genome Relatedness Index) estimations and phylogenomic analyses performed through the Type Strain Genome Server (TYGS) (Meier-Kolthoff and Göker, 2019) and TrueBac ID (Ha *et al*., 2019) services. Average Nucleotide Identity (ANI) between GeG2^T^ genome and 7,184 Alphaproteobacteria genomes downloaded from NCBI (Accessed in July 2019) was calculated with FastANI (Jain et al., 2018), using default parameters.

Whole genome alignment between the strain GeG2^T^ genome and *N. rosa* NBRC15208 (GCF_001598555.1), the only genome available for the species in NCBI, was performed with MUMmer toolkit v3.1 (Kurtz et al., 2004). Alignments with at least 1 kb length and 90% identity were used to draw a circular visualization of alignments with ggbio (Yin et al., 2012).

### Genome functional annotation and predictions

Genome annotation was performed with Prokka v1.14.0 (Seemann, 2014) and rRNA sequences were predicted with Barrnap v0.9 (github.com/tseemann/barrnap). Gene functions based on KEGG Orthology were predicted with <u>KofamScan</u> v1.3.0 (Aramaki et al., 2019), all orchestrated with the bacannot pipeline (github.com/fmalmeida/bacannot). Plasmid sequences were predicted using Plasflow v1.1 (Krawczyk et al. 2018) and sequence similarity against known plasmids was calculated with the NCBI’s Microbial Nucleotide Blast. Dinucleotide relative abundance distance between genome replicons were determined as described in diCenzo and Finan (2017). Clusters of Orthologous Genes (COG) assignments were performed by eggNOG-mapper v2 (Huerta-Cepas et al. 2017) and the Two-Proportions Z-Test was used with the *prop.test,* R language built-in function, to calculate significance between the proportions of COG categories annotated in the replicons. Metabolic pathways and oxygenases were determined with KEGG Orthology database (Kanehisa et al., 2008),

## RESULTS AND DISCUSSION

### Isolation, growth and 16S rDNA based phylogenetic analysis

A bacterial strain, denominated GeG2^T^, was isolated from microbial enrichments obtained from Cerrado soils, an extremely biodiverse environment with vast microbial genetic repertoire and biotechnological potential (Alves-Prado et al., 2010; Peixoto et al., 2017). Growth in solid minimal medium (MM), resulted in white, circular (2-3 mm diameter), convex colonies, with regular edges and shiny appearance. Light microscopy analysis revealed GeG2^T^ cells to be Gram-negative, motile, rod-shaped, with dimensions ranging from 1.3–2.3 μm length and 0.3–0.6 μm width.

Nearly complete 16S rRNA gene fragments of strain GeG2^T^ were obtained by PCR amplification and sequenced. All 22 sampled amplicon sequences (1,450 bp) were identical and comparisons with sequences from EZBioCloud database (Yoon et al., 2017) revealed the highest similarity with *Novosphingobium rosa* NBRC15208 (100% rDNA sequence identity, with 95% of sequence coverage), followed by *Novosphingobium lotistagni* THG-DN6.20 (97.58%), *Novosphingobium oryzae* ZYY112 (97.02%) and *Novosphingobium barchaimii* LL02 (96.95%). Further phylogenetic reconstructions based on 16S rRNA genes also evidenced the placement of strain GeG2^T^ in the alphaproteobacterial genus *Novosphingobium* and its close relatedness with the species *N. rosa* (Supplementary Figure S1). Although highly useful for initial taxonomic classification of microorganisms, resolution limitations in phylogenetic analysis based solely on 16S RNA are often reported, especially at the species level (Fox et al., 1992; Stackebrandt and Goebel, 1994; Jaspers and Overmann, 2004), indicating that additional approaches must be employed for the delimitation of bacterial species (Chun et al., 2018; Raina et al., 2019). For this reason, physiological, chemical, morphological and whole genome-based analyses of strain GeG2^T^ were carried out, as described in the following sections.

### Physiological and chemotaxonomic characterizations

Considering the highest 16S rRNA gene identity observed between strain GeG2^T^ and *Novosphingobium rosa*, a type strain of this species, DSM7285^T^, was obtained from DSMZ microorganism collection (Leibniz Institute, Germany) and used as the reference strain for comparative phenotypic analyses. Differently from *N. rosa* DSM7285^T^, strain GeG2^T^ was unable to grow in several rich media, such as Trypticase Soy Agar, Nutrient Agar and Luria-Bertani medium (Table 1). Some *Sphingomonadaceae* members isolated from soils are known to exhibit better growth in low nutrient culture media (Gupta et al., 2009; Glaeser and Kämpfer, 2014) and the characteristic low nutrient availability of Cerrado soils (Haridassam, 2008) could probably have driven GeG2^T^ strain adaptation to oligotrophic growth conditions.

**Table 1.** Differential characteristics of strain GeG2^T^ and type strains of related *Novosphingobium* species. Strains: 1. GeG2^T^; 2. *N. rosa* DSM7285^T^ (data from this study); 3. *N. lotistagni* THG-DN6.20^T^ (data from Ngo et al. 2016). All strains are positive for hydrolysis of aesculin; catalase; oxidase; assimilation of glucose, arabinose and maltose; alcaline phosphatase, acid phosphatase, naphtol-AS-BI- phosphohydrolase, β-galactosidase, α-glucosidase and β-glucosidase activities. All strains are negative for hydrolysis of gelatin and urea; indol production; glucose fermentation; assimilation of mannitol, caprate, adipate, malate, citrate, phenylacetate; arginine dihydrolase, lipase (C14) and α-mannosidase activities. +, positive; −, negative; w, weak positive; -* no growth after first transfer; ND, no data.

| Characteristic | 1 | 2 | 3 |
| --- | --- | --- | --- |
| Isolation source | Cerrado soil | Rose rhizosphere | Lotus pond |
| Colony color | White | Yellow | Yellow |
| Cell size (μm): | | | |
| Width | 0.3 - 0.6 | 0.3 - 0.5 | 0.6 - 0.8 |
| Length | 1.3 - 2.3 | 1.0 - 1.9 | 0.9 - 4.2 |
| Motility | + | + | - |
| Growth on TSA | - | + | w |
| Growth on NA | -* | + | + |
| Growth on LB | -* | + | - |
| Growth conditions: |  |  |  |
| pH | 4-7 | 4-7 | ND |
| NaCl (%) | 0-1 | 0-1.5 | 0-1 |
| Temperature ($^{\circ}$C) | 15-33 | 15-33 | 4-42 |
| Facultatively anaerobic growth | + | - | - |
| Hydrolysis of starch | + | - | - |
| Nitrate reduction | w | - | - |
| **Assimilation of:** |  |  |  |
| Mannose | + | + | - |
| N-Acetylglucosamin | + | w | w |
| Gluconate | + | w | - |
| **Enzyme activities of:** |  |  |  |
| Esterase (C4) | w | + | w |
| Esterase Lipase (C8) | w | w | w |
| Leucin-Arylamidase | + | + | w |
| Valin-Arylamidase | + | + | w |
| Cystin-Arylamidase | w | w | w |
| Trypsin | - | - | + |
| Chymotrypsin | w | - | + |
| α-Galactosidase | - | - | + |
| β-Glucuronidase | w | - | + |
| N-Acetyl-β-Glucosaminidase | - | w | - |
| α-Fucosidase | - | - | w |
| DNA G+C content (mol%) | 63.57 | 64.50 | 63.10 |

Morphological differences were observed between colonies of strain GeG2^T^ and *N. rosa* DSM7285^T^ grown in MM for 48 hours, with colonies of strain GeG2^T^ presenting a whitish and viscous aspect, while *N. rosa* DSM7285^T^ colonies were smaller, drier and yellow-pigmented (Supplementary Figure S2). Both bacterial strains grow in temperatures between 15 to 33 °C, with optimal growth at 28 °C, and pH 4.0 to 7.0, though while *N. rosa* DSM7285^T^ grows in NaCl concentrations ranging from 0 to 1.5%, strain GeG2^T^ only grows up to 1% NaCl (Table 1). Starch hydrolysis was not observed for *N. rosa* DSM7285^T^, whereas GeG2^T^ was positive after 14 days of incubation in MM containing both glucose and starch, or only starch (Table 1, Supplementary Figure S3). Furthermore, even though *Novosphingobium* members were initially described as strict aerobes (Takeuchi et al., 2001), some species have recently been identified as facultative anaerobes (Sohn et al., 2004; Sheu et al., 2020). As reported for *N. pentaromativorans* US6-1^T^ (Sohn *et al*., 2004), the slow growth of strain GeG2^T^ under anaerobic conditions was detected (after 30 days), which was never observed for *N. rosa* DSM7285^T^ (Table 1).

Antibiotic susceptibility profiles based on disc-diffusion tests (see Methods) revealed strain GeG2^T^ to be resistant to several antibiotics, with susceptibility observed only to tetracycline and rifampicin. A similar resistance pattern was observed for *N. rosa* DSM7285^T^, which, in addition to tetracycline and rifampicin, was also susceptible to trimethoprim/sulfamethoxazole. Although information regarding resistance mechanisms in *Sphingomonadaceae* is still scarce, soil and rhizosphere isolates belonging to this family are generally resistant to various kinds of antimicrobial agents (Yabuuchi et al., 2002) and have been identified to constitute an environmental reservoir of the antibiotic resistome (Dantas et al., 2008).

Additional biochemical characterizations were performed with API 20NE and API Zym kits and revealed strain GeG2^T^ to be positive for aesculin hydrolysis, catalase and oxidase activity, assimilation of glucose, arabinose, mannose, N-acetyl-glucosamine, maltose, gluconate and enzymatic activities of alkaline phosphatase, leucin-arylamidase, valin-arylamidase, acid phosphatase, naphtol- AS-BI-phosphohydrolase, β-galactosidase, α-glucosidase and β-glucosidase. Furthermore, weak positive activities were observed for esterase, esterase lipase, cystin-arylamidase, chymotrypsin and β- glucuronidase, as well as for nitrate reduction. Similar overall biochemical profiles were observed for *N. rosa* DSM7285^T^, although it was negative for nitrate reduction, chymotrypsin and β-glucuronidase activities and weak positive for N-acetyl-β-glucosaminidase.

As expected for members of *Sphingomonadaceae* family (Takeuchi et al., 2001; Glaeser and Kämpfer, 2014), the major respiratory quinone identified in GeG2^T^ cells was Q10 (>95%), with Q9 also detected in smaller amounts. Major fatty acids were C16:0 (24.62%), C18:0 (21.84%) and C18:1ω7ϲ/ C18:1ω6ϲ (18.18%) (Table 2). Despite presenting similar composition, proportions of different fatty acids varied considerably between GeG2^T^ and *N. rosa* DSM7285^T^, for which the major fatty acids detected were C18:1ω7ϲ/ C18:1ω6ϲ (38.71%), followed by C14:0 2-OH (19.65%) and C16:0 (14.84%) (Table 2). Notably, while the unsaturated fatty acid C18:1ω9ϲ represented a considerable fraction of fatty acids (8.29%) in GeG2^T^ cells, it was only detected in trace amounts (<1%) in *N. rosa* DSM7285^T^ cells grown under the same conditions (Table 2).

**Table 2.** Whole-cell fatty acid contents (%) of strain GeG2^T^ and the closest related species *Novosphingobium rosa* DSM7285 ^T^. Both strains were grown on MM agar at 28 °C for 72 hours. TR, Trace (<1%); -, not detected.

| Fatty acid | GeG2<sup>T</sup> | *N. rosa* DSM7285<sup>T</sup> |
| --- | --- | --- |
| C<sub>12:0</sub> | TR | - |
| C<sub>13:0</sub> anteiso | TR | TR |
| C<sub>14:0</sub> | 1.85 | 1.30 |
| C<sub>14:0</sub> 2-OH | 7.22 | 19.65 |
| C<sub>16:0</sub> | 24.62 | 14.84 |
| C<sub>17:0</sub> anteiso | TR | - |
| C<sub>17:0</sub> cyclo | TR | - |
| C<sub>17:0</sub> | TR | - |
| C<sub>18:1</sub> ω9c | 8.29 | TR |
| C<sub>18:0</sub> | 21.84 | 7.82 |
| C<sub>19:0</sub> iso | TR | - |
| C<sub>19:0</sub> cyclo ω8c | 1.92 | 2.18 |
| C<sub>20:0</sub> | TR | - |
| C<sub>16:1</sub> ω7c/C<sub>16:1</sub> ω6c | 11.97 | 13.88 |
| C<sub>18:1</sub> ω7c/ C<sub>18:1</sub> ω6c | 18.18 | 38.71 |

The polar lipids of strain GeG2^T^ included phosphatidylglycerol (PG), phosphatidylethanolamine (PE), sphingoglycolipid (SGL), diphosphatidylglycerol (DPG), besides an aminolipid (AL) and unidentified lipids (Supplementary Figure S4). Interestingly, even though phosphatidylmonomethylethanolamine (PME) and phosphatidyldimethylethanolamine (PDE) are commonly identified in *Novosphingobium* members (Glaeser *et al*., 2013), including *N. rosa* (Kämpfer *et al*., 1997), these polar lipids were not detected in strain GeG2^T^. Differences in polar lipids and fatty acids profiles between strain GeG2^T^ and the *N. rosa* strain suggest that these bacteria do not belong to the same species, as sphingomonads species can be distinguished from each other due to qualitative and/or quantitative variations in these chemotaxonomic components (Busse et al., 1999).

### Protein profile analyses by MALDI-TOF

Comparative analysis of protein profiles by MALDI-TOF mass spectrometry was performed to further demonstrate the distinctive phenotype of strain GeG2^T^. While main spectra profiles (MSP) obtained for strain DSM7285^T^ matched those from the species *Novosphingobium rosa* (scores >2.0), MSP generated from GeG2^T^ protein extracts showed identification scores lower than 2.0 when compared to DSM7285^T^ MSP and MALDI Biotyper® database, suggesting that this strain represents a new species, not yet represented in the database containing over 8,000 strains. Furthermore, dendrogram analysis of generated MSP shows that GeG2^T^ and *N. rosa* DSM7285^T^ are grouped in different clades, even when external groups are added (Supplementary Figure S5), further indicating that they belong to different species.

### Morphological characterization of GeG2^T^ cells by electronic microscopy

Transmission electron micrographs of strain GeG2^T^ cells revealed one or two intracytoplasmic electron-dense granules (70–150 μm diameter) per cell, mostly located in the central portion of the cytoplasm (Figures 1A and 1B). The observed granules present a characteristic aspect of polyphosphate (poly-P) granules or acidocalcisomes, subcellular structures enriched in phosphorus compounds and different cations (Docampo, 2006). While acidocalcisomes are membrane-encapsulated, poly-P granules lack surrounding membranes and, even though recent studies suggested that these are different subcellular structures that can be simultaneously found in alphaproteobacterial species (Frank and Jendrossek, 2020), the presence of surrounding membranes in GeG2^T^ granules could not be clearly determined by electron microscopy. Further microscopic and chemical analyses are required to elucidate the nature and function of the electron-dense granules observed in strain GeG2^T^.

**Figure 1.**
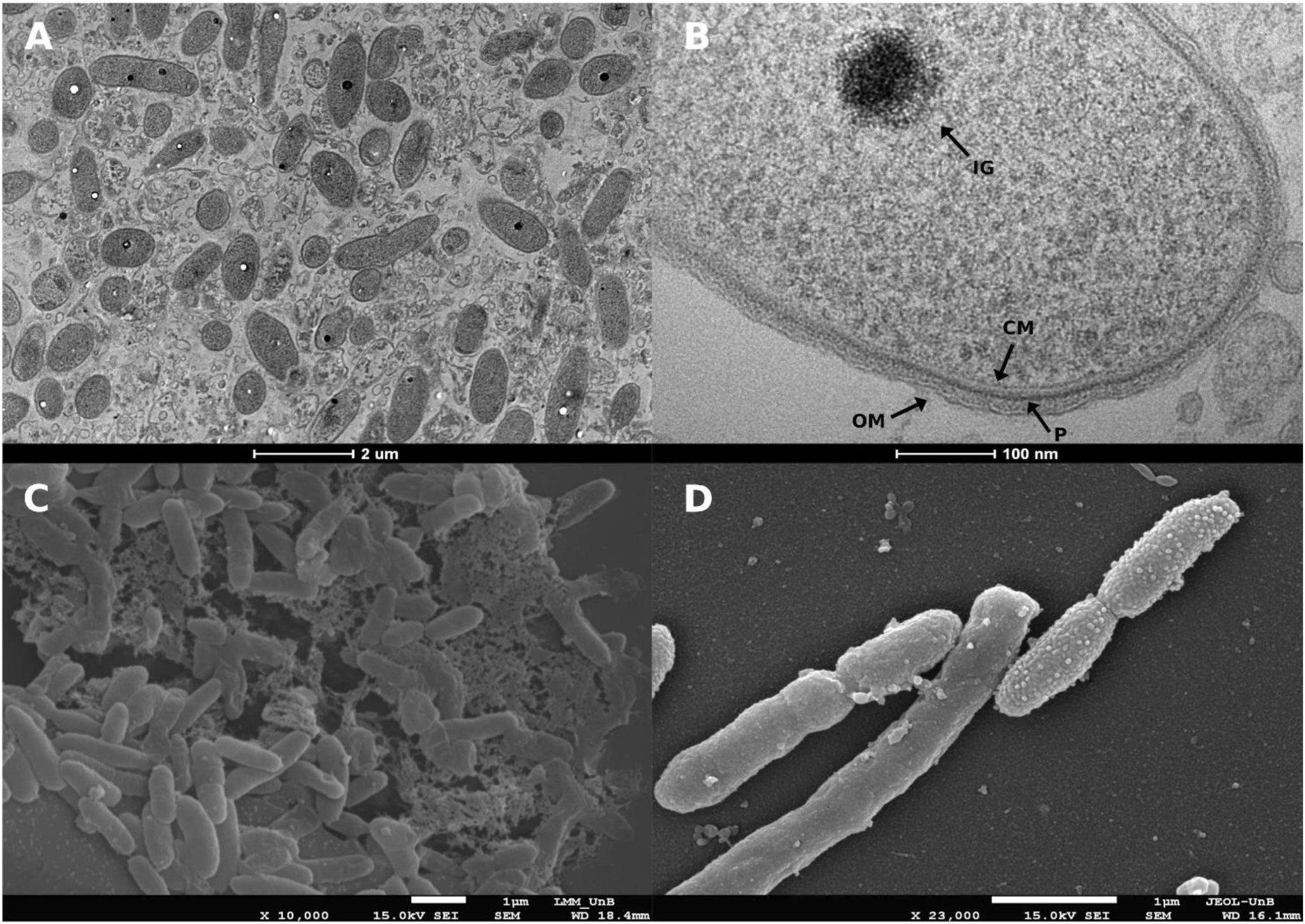
Transmission electron micrograph of strain GeG2^T^ cells showing intracytoplasmic electron- dense granules (**A** and **B**) and scanning electron micrographs of strain GeG2^T^ cells surrounded by amorphous polymeric matrix, with characteristic aspect of exopolysaccharides (EPS) (**C**) and presenting smooth or granular surfaces (**D**). Magnifications and scale bars are indicated under each micrograph. IG: Intracytoplasmic granule. CM: Cytoplasmatic membrane. OM: Outer membrane. P: Peptidoglycan layer.

Scanning electron micrographs of GeG2^T^ cells grown in liquid MM revealed aggregated cells surrounded by an amorphous polymeric matrix, with characteristic aspect of exopolysaccharides (EPS) (Figure 1C). *Sphingomonadaceae* members are known for their ability to produce diverse EPS, generically denominated sphingans, presenting important applications in bioremediation, pharmaceutical and food industries (Wu et al., 2017). Moreover, heterogeneities in cell surfaces among different GeG2^T^ cells have also been identified: some showing smooth uniform aspect, whilst others were granular and rough, a characteristic aspect of outer membrane vesicle (OMV) producers (Figure 1D) (Gilewicz *et al*., 1997; Coppotelli *et al*., 2010). Although initially characterized in pathogenic Gram-negative bacteria (Toyofuku et al., 2019), OMVs produced by environmental species have been increasingly studied and their varied composition under different conditions suggest diverse biological functions, including nutrient acquisition, stress responses, biodegradation of aromatic compounds, surfactant actions and biofilm formation (Coppotelli et al., 2010; Choi et al., 2014; Schwechheimer and Kuehn, 2015; Yun et al., 2017; De Lise et al., 2019).

### Planktonic-sessile dimorphism of strain GeG2^T^

After sequential transfers in liquid MM, isolate GeG2^T^ was found to grow as either planktonic motile cells or sessile-aggregated non-motile cells that form macroscopic flocks of different sizes (Supplementary Figure S6). This phenomenon, also observed in other sphingomonads, is known as planktonic sessile dimorphism (Pollock and Armentrout, 1999, Tiirola *et al*., 2002; Notomista *et al*., 2011). As reported for other species (Pollock and Armentrout, 1999), flock formation by strain GeG2^T^ seemed to be influenced by different growth parameters. While culture agitation increased flock formation, cultures incubated without agitation or in flasks containing greater volumes of media showed fewer macroscopic flocks and more homogeneous turbidity, indicating that greater oxygen levels may favor cell aggregation and extracellular matrix production. Increased flock formation was also observed in non-agitated cultures grown for extended incubation periods (over four days).

Scanning electron micrographs from agitated cultures grown for 14 days and presenting several flocks revealed long extracellular fibers forming a network connecting the cells, as well as extracellular matrix enclosing them (Figure 2A). An increased number of cells with granular surfaces were also identified, suggesting that OMV production could be favored in these cultivation conditions (Figure 2B). Moreover, spherical structures resembling large vesicles, with diameters ranging from 150 to 250 nm, were identified among cell aggregates (Figure 2B). Similar extracellular structures were reported for environmental bacteria grown in minimal media containing different hydrocarbons, in which an emulsifying activity and optimization of substrate assimilation roles were proposed (Goutx *et al*., 1987; Husain *et al*., 1997). Furthermore, formation of macroscopic flocks composed of aggregated cells embedded in extracellular matrix rich in size-varying vesicle structures have recently been described for *Novosphingobium* sp. PP1Y grown in minimal media containing glutamate as sole carbon source, suggesting a potential role of vesicles in nutrient acquisition under limiting conditions (De Lise *et al*, 2019). Likewise, it is possible that the vesicles associated with GeG2^T^ cell aggregates observed after long incubation periods may be related to better nutrient assimilation under oligotrophic conditions.

**Figure 2.**
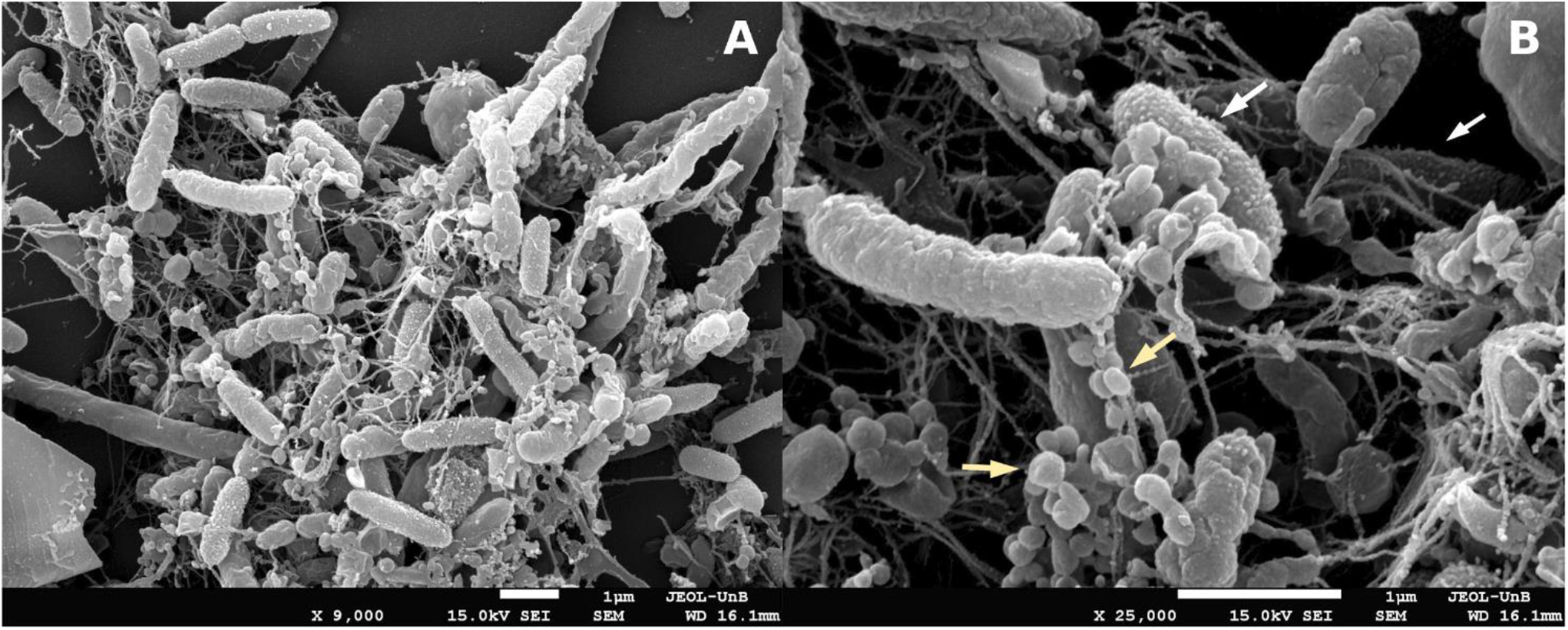
Scanning electron micrographs of strain GeG2^T^ cultures presenting macroscopic flocks grown in MM for 14 days. Cells presenting granular surfaces are indicated by white arrows and spherical structures resembling large vesicles are indicated by yellow arrows. Magnifications and scale bars are indicated under each micrograph.

### Genome assembly and genome-based taxonomic analyses

To further improve its characterization, the complete genome sequence of strain GeG2^T^ was assembled using a combination of long and short-reads sequencing approaches, resulting in a high-quality and contiguous assembly totalizing 7,162,928 bp, distributed in 6 contigs, with a total GC content of 63.57% (Table 3). Four circular replicons were identified: a 4,164,843 bp chromosome (GC content: 64.23%), a 2,710,928 bp extrachromosomal megareplicon (GC content: 62.93%) and two plasmids – pGeG2a and pGeG2b - with 212,687 and 68,553 bp (GC content of 61.41 and 55.61%, respectively) (Figure 3). Two additional contigs containing the entire rRNA operon and tRNA genes (with 5,558 bp and 359 bp) were assembled separately from the chromosome by Unicycler (Supplementary Figure S7) possibly due to difficulties associated with the resolution of highly repetitive regions (Wick *et al*., 2017).

**Figure 3.**
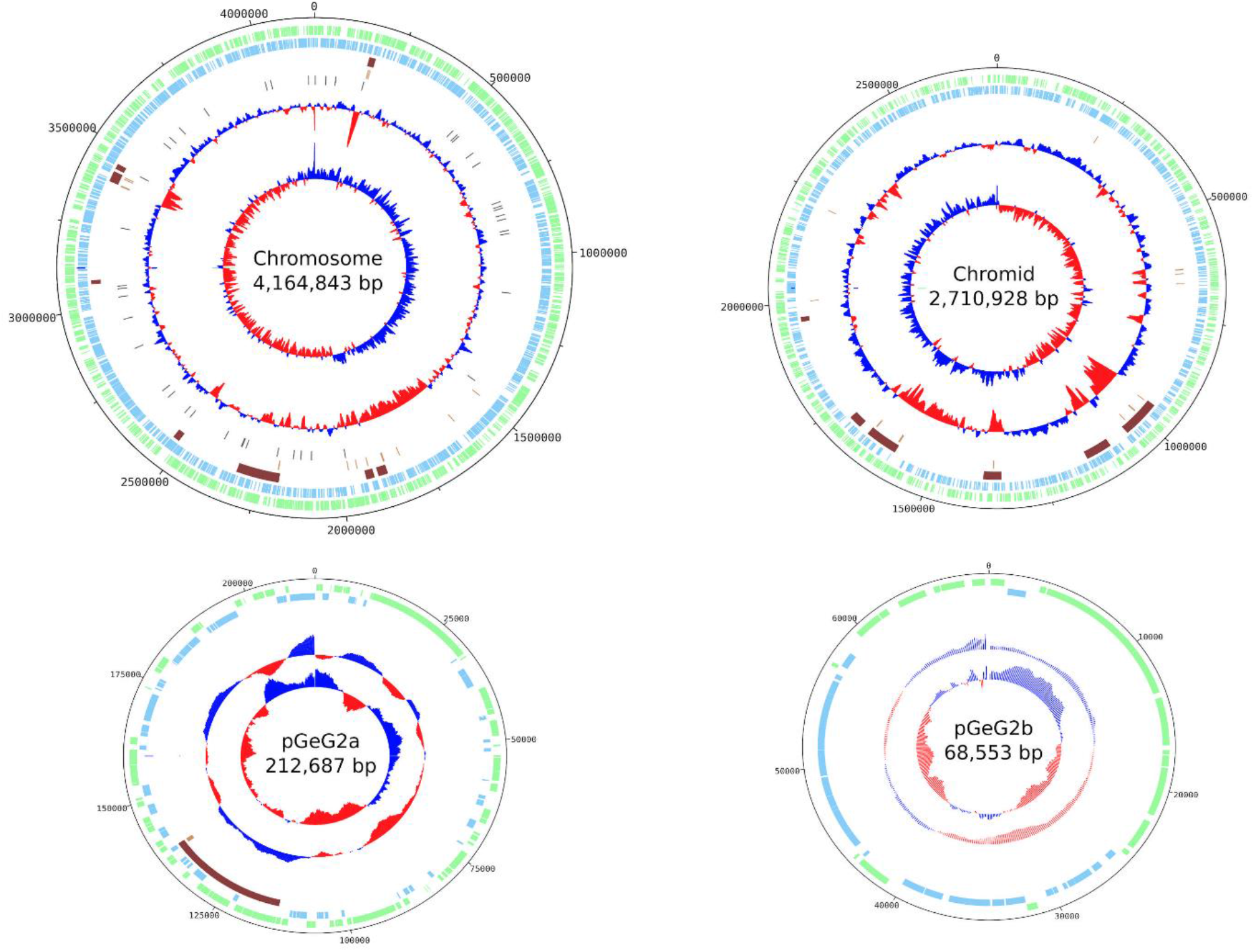
Circular maps and genetic features of the chromosome, chromid and plasmids of *Novosphingobium* sp. strain GeG2^T^. From outside to center: forward CDS (green), reverse CDS (blue), genomic islands (brown), transposases (pink), tRNAs (purple), GC content and GC skew (red and blue). Replicons are not shown to scale.

**Table 3.** Statistics of strain GeG2^T^ genome assembly performed with a hybrid approach using both short (Illumina) and long reads(PacBio).

| Feature | Value |
| --- | --- |
| Number of contigs | 6 |
| Largest contig (bp) | 4,164,843 |
| Total size (bp) | 7,162,928 |
| N50 | 4,164,843 |
| L50 | 1 |
| GC (%) | 63.57 |

The complete genome sequence of strain GeG2^T^ offers a new perspective regarding its relationship with *N. rosa*, given the context of 100% sequence identity of their 16S rRNA genes. Initially, we could ascertain that GeG2^T^ 16S rRNA gene sequence predicted from the genome assembly (1,486 bp) was identical to the sequences obtained by direct colony PCR amplification followed by Sanger sequencing (1,450 bp). As recently proposed by Chun et al. (2018), overall genome related indexes (OGRI) were calculated in order to better evaluate the taxonomic classification of strain GeG2^T^. As shown in Figure 4, Average Nucleotide Identity (ANI) and digital DNA:DNA hybridization (dDDH) values obtained for these isolates (90.38% and 42.60%) are markedly divergent, considering the generally accepted species boundary of 95-96% and 70% for ANI and dDDH, respectively (Chun et al., 2018). Moreover, genome-based taxonomic analyses performed in both TYGS and TrueBac ID servers identified that strain GeG2^T^ does not belong to any species currently found in their databases, further indicating that it represents a new species within the genus *Novosphingobium* (Supplementary Figure S8, Supplementary File S1).

**Figure 4.**
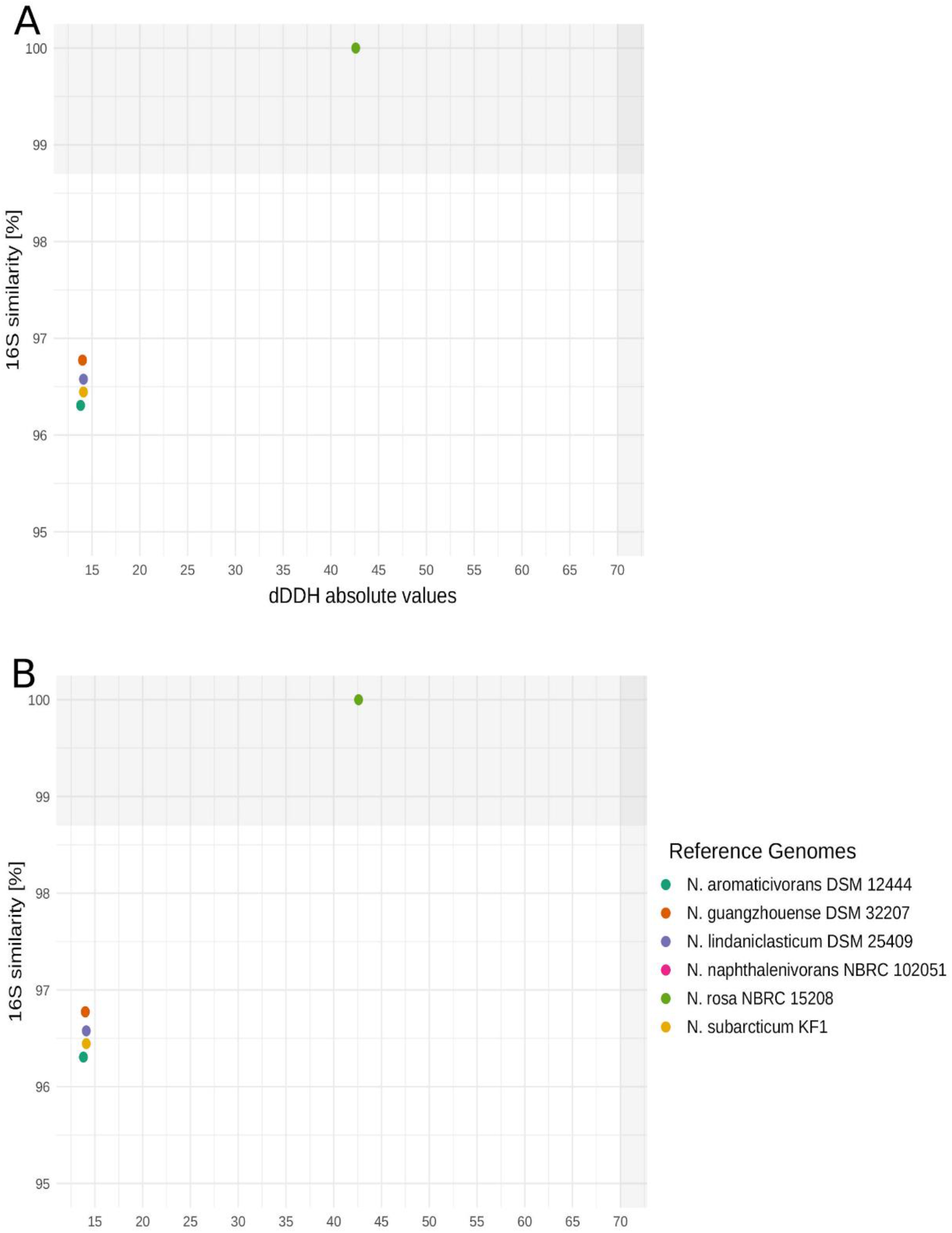
Relationship between 16S rRNA gene sequence similarities and ANI values (**A**) or dDDH values (**B**) for *Novosphingobium* sp. GeG2^T^ and six genome-sequenced type strains in the genus *Novosphingobium*. The species boundary of ANI and dDDH values are indicated at 95 and 70%, respectively (Chun et al., 2018).

Whole genome alignment between the strain GeG2T and *N. rosa* NBRC15208 indicates that the genomes are divergent from each other. When setting an alignment threshold of 90% identity in blocks larger than 1 kb, only 77% of the bases in the chromosome and 38% in the megareplicon could be mapped. A circular visualization of the whole genome alignment between these genomes is shown in Figure 5 and allows the observation of several sizeable unaligned regions, highlighting the dissimilarity between the genomes.

**Figure 5.**
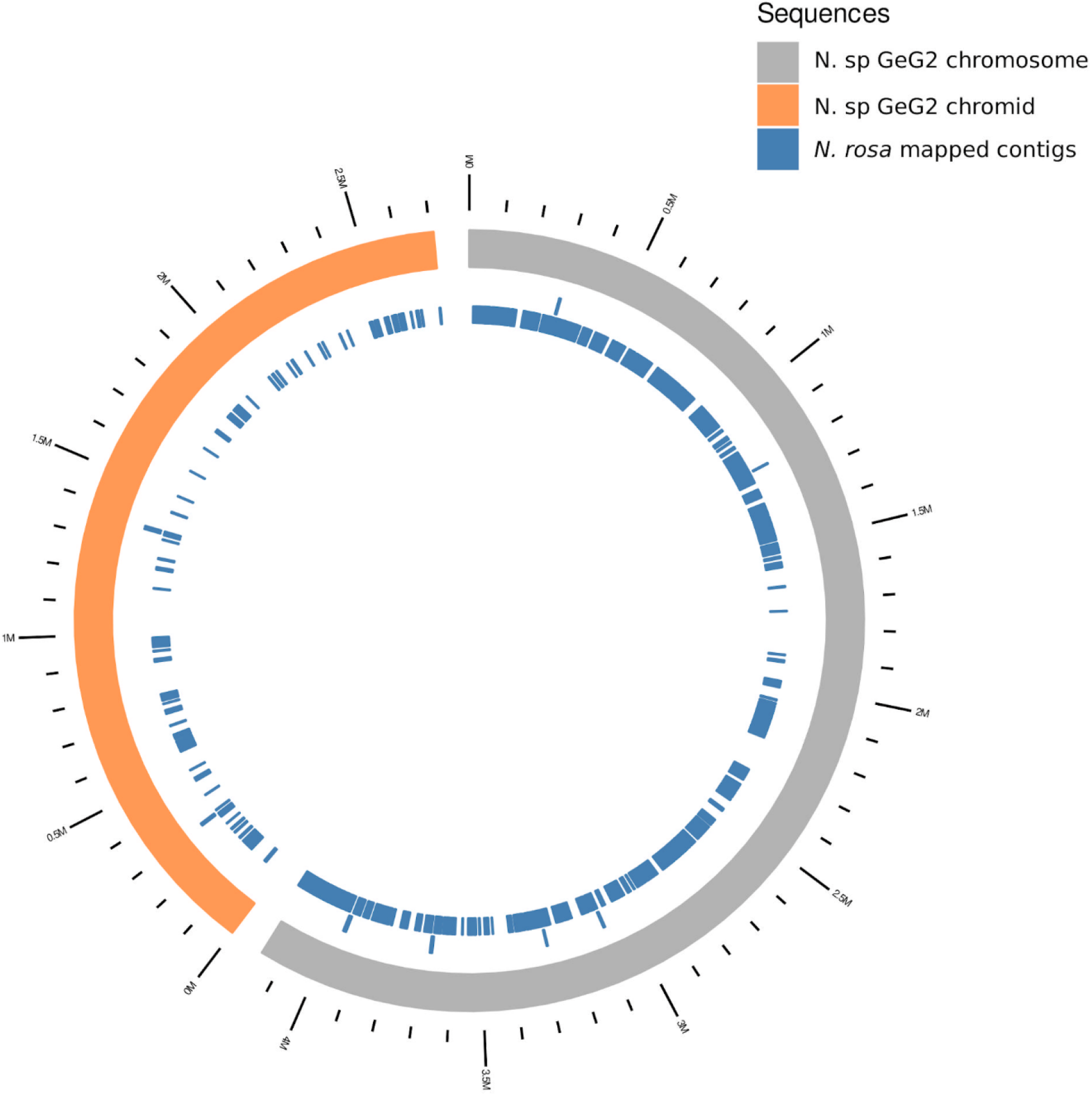
Circular visualization of whole genome alignment between the strain GeG2^T^ (chromosome in grey and chromid in orange) and *N. rosa* NBRC15208 reference genome (in blue). Genome alignments have been filtered to minimum length of 1,000 nucleotides and identity of 90%.

### The genome is bipartite and contains a chromid

Besides the chromosome and two plasmids, an interesting feature identified in the genome of strain GeG2^T^ was the presence of an extrachromosomal megareplicon (2.7 Mb) (Figure 3). Gene content and genomic signature analyses of this replicon revealed both chromosomal and plasmid features, commonly associated with chromids: replicons normally larger than the accompanying plasmids but smaller than the chromosome, presenting nucleotide composition close to that of the chromosome, but plasmid-type replication and maintenance systems (Harrison et al., 2010).

While plasmid replication and segregation genes belonging to the alphaproteobacterial RepABC family have been identified in the megareplicon of strain GeG2^T^, close GC content (Δ=1.2%) and dinucleotide relative abundance distance from the chromosome (<0.4) were observed, supporting its classification as a putative chromid according to diCenzo and Finan (2017). Furthermore, the presence of one or more core genes that can be found in the chromosome of related species is considered a distinctive characteristic of chromids (Harrison et al., 2010). Indeed, genes involved in important cellular functions, such as DNA polymerase III ε and α subunits (genes *dnaQ* and *dnaE*), pentose phosphate and pyruvate oxidation pathway genes were detected both in the chromosome and in the putative chromid of strain GeG2^T^, based on KEGG annotations. Moreover, the alignment of sequences with coding potential predicted in the chromosome and the megareplicon has allowed the identification of 29 genes with high nucleotide and amino acid sequence similarities (>85%) (Supplementary File S2).

Secondary megareplicons (>350 kb) were identified in all six fully resolved *Novosphingobium* genomes available in databases so far, suggesting that multipartite genomes may be a common characteristic of this genus (Wang et al., 2018; Gogoleva et al., 2019). The putative chromid identified in the genome of strain GeG2^T^ is the largest secondary replicon reported for *Novosphingobium* species to date, with 2.7 Mb, followed by a 2.2 Mb replicon identified in *Novosphingobium* sp. P6W (Gogoleva et al., 2019), a 1.7 Mb replicon from *N. resinivorum* SA1 (Wang et al., 2018) up to a 487.3 kb putative chromid in the genome of *N. aromaticivorans* DSM12444 (diCenzo and Finan, 2017). Interestingly, the *N. rosa* NBRC15208 genome assembly (GCF_001598555.1) used for comparative purposes with GeG2^T^ does not report any plasmids or chromids. However, the pairwise genome alignment, shown in Figure 5, reveals several NBRC15208 contigs mapping to GeG2^T^ chromid, indicating that *N. rosa* NBRC15208 might have a chromid that could not be resolved due to the exclusive use of short-reads for this genome assembly. This underscores the benefits of hybrid assembly using both short and long reads underlying the high-quality assembly obtained for GeG2^T^ and suggest that megareplicons could be more widespread in *Novosphingobium* genomes than currently acknowledged.

Next, we set out to explore if the chromosome and chromid of GeG2^T^ share sequence identity enabling to trace their evolutionary origins. In order to investigate such hypothesis, DNA sequence alignments of both replicons were performed using blastn and results were filtered to keep only hits with 100% similarity and longer than 1,000 bp. Using these thresholds, only six hits of approximately 1,300 bp have been identified. Interestingly, we found a single region of the chromid (between the positions 1,657,174 and 1,655,854) matching three different locations in the chromosome. Inside this genomic region, two IS3 family transposases, namely IS868 and ISRtr2, have been found *in tandem*. Altogether, these results indicate distinct evolutionary routes between the chromosome and chromid, and that transposable elements may mediate DNA exchange between these megareplicons in GeG2^T^, even though in a very limited scale.

### Gene annotation

In total, 6,124 genes (3,697 in the chromosome) were predicted in the genome of strain GeG2^T^, including 3 rRNAs (collapsed in the rRNA operon in contig 5), 57 tRNAs, 1 tmRNA and 6,063 protein coding sequences (CDS). A putative function was assigned to 3,706 of the protein-coding genes, while 2,357 were annotated as hypothetical proteins.

Functional analysis based on clusters of orthologous groups of proteins (COGs) assigned 5,746 CDS predicted from strain GeG2^T^ genome to one or more COG functional classes, which were then sorted into 21 groups, as shown in Figure 6A. As frequently observed in bacterial multipartite genomes (Mackenzie et al., 2001; Chain et al., 2006; Jansen et al., 2010), functional biases were detected between the genomic replicons of strain GeG2^T^. While proteins involved in essential processes such as DNA replication and repair (L), cell division (D), translation machinery (J) and cell wall/ membrane biogenesis (M) are mostly present in the chromosome, an enrichment of genes involved in inorganic ion transport and metabolism (P), transcription (K) and carbohydrate transport and metabolism (G) is observed in the secondary megareplicon (Figure 6A). These categories were shown to be primarily enriched in chromids rather than in megaplasmids (DiCenzo and Finan, 2017), further corroborating the classification of GeG2^T^ megareplicon as a chromid and indicating that it is likely related to adaptative roles and responsive behaviors to environmental changes (Jansen et al., 2010). However, unlike other bacterial species in which cellular motility (N) and signal transduction mechanisms (T) categories are overrepresented in chromids (Jansen *et al*., 2010; Frank *et al*., 2015), most genes associated with these functions are chromosomal in strain GeG2^T^ (Figure 6A). Also, the majority of putative proteins assigned to the unknown function COG category (S) are present in the chromosome, differently from others sphingomonads, in which hypothetical proteins were mainly found in secondary replicons (Aylward et al., 2013).

**Figure 6.**
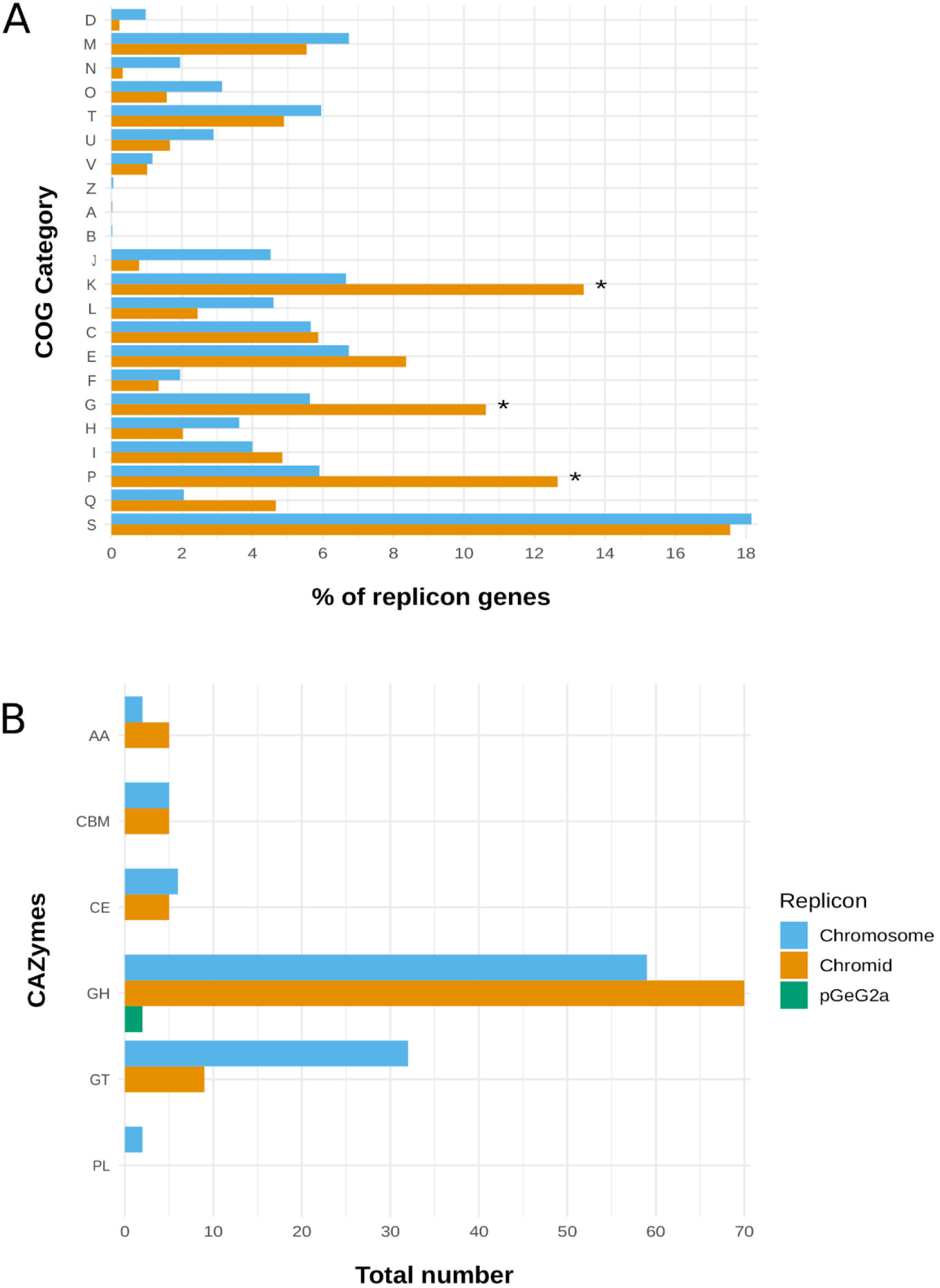
COG functional categories (**A**) and carbohydrate-active enzymes (CAZymes) (**B**) predicted in each replicon of strain GeG2^T^ genome. D: Cell cycle control, cell division, chromosome partitioning; M: Cell wall/membrane/envelop biogenesis; N: Cell motility; O: Post-translational modification, protein turnover, chaperone functions; T: Signal transduction mechanisms; U: Intracellular trafficking, secretion and vesicular transport; V: Defense mechanisms; Z: Cytoskeleton; A: RNA processing and modification; B: Chromatin structure and dynamics; J: Translation, ribosomal structure and biogenesis; K: Transcription; L: Replication and repair; C: Energy production and conversion; E: Amino Acid metabolism and transport; F: Nucleotide metabolism and transport; G: Carbohydrate metabolism and transport; H: Coenzyme metabolism and transport; I: Lipid metabolism and transport; P: Inorganic ion transport and metabolism; Q: Secondary metabolites biosynthesis, transport and catabolism; S: Function unknown. GT: glycosyl transferases; GH: glycosyl hydrolases; CE: carbohydrate esterases; CBM: carbohydrate binding module; AA: enzymes for the auxiliary activities; PL: polysaccharide lyases.

Regarding the plasmids detected in strain GeG2^T^ (pGeG2a and pGeG2b), sequence similarity searches against the NCBI database did not result in any significant alignment, indicating they are newly identified plasmids with still unknown functional properties, as depicted by the fact that most CDS detected in these replicons were annotated as hypothetical proteins. Genes related to cation efflux (*cusC*) and drug efflux systems (*emrB, emrK, stp*) have been predicted in pGeG2a, suggesting a potential role of this plasmid in drug resistance (Li and Nikaido, 2009). Moreover, functional analyses based on COGs revealed that while transcription (K), amino acid transport/metabolism (E) and carbohydrates transport/metabolism (G) were the main functional categories detected in pGeG2a, proteins predicted in pGeG2b were predominantly (almost 40%) attributed to trafficking, secretion and vesicular transport category (U) (Supplementary Table S1), frequently associated with resistance to toxic compounds (Zheng et al., 2015; diCenzo e Finan, 2017; Suzuki et al., 2019). Considering the indication of vesicle production by strain GeG2^T^ detected in SEM micrographs (Figure 2B), it is tempting to speculate that the numerous genes from this functional category identified in its plasmid pGeG2b, as well as in its chromosome and chromid (Figure 6A), could be related to this phenotype. Furthermore, a considerable fraction of predicted proteins in both plasmids were also attributed to replication, recombination and repair functions (category L) (Supplementary Table S1). Gene enrichments from categories L and K have been reported for plasmids of several bacterial species and are possibly associated with replication and conjugation processes (Zheng et al., 2015; diCenzo and Finan, 2017) and the identification of tra and trb genes in both plasmids from strain GeG2^T^ indicates their conjugative nature.

### Biochemical pathways inferred from gene annotation

Strain GeG2^T^ genome encodes an extensive repertoire of enzymes involved in carbohydrate metabolism, comprising 41 glycosyl transferases (GTs), 131 glycosyl hydrolases (GHs), 11 carbohydrate esterases (CEs), two polysaccharide lyases (PLs) and seven enzymes for auxiliary activities (AAs), totalizing 192 carbohydrate-active enzymes (CAZymes; Figure 6B, Supplementary File S3), a higher number than previously reported for other *Sphingomonadaceae* (Aylward et al., 2013; D’Argenio et al., 2014). As shown by COG analysis, many genes encoding CAZymes are found in the chromid (89; 47%), especially GHs and AAs (Figure 6B), indicating an important role of this replicon in polysaccharide catabolism (Nguyen et al., 2018).

Among GHs encoded in the genome, 40 (23 in the chromid and 17 in the chromosome) belonging to families related to the degradation of xylan and other hemicellulose components (GH3, GH5, GH8, GH10, GH16, GH30, GH43, GH51, GH67, GH115) were identified, as well as CEs involved in the removal of the side chains from substituted xylose units (CE1 and CE4), revealing a large potential of strain GeG2^T^ for plant biomass degradation (Nguyen et al., 2018). Moreover, five GHs classified in families GH106 and GH78 of α-L-rhamnosidases, enzymes involved in many essential microbial functions and biotechnological applications (Manzanares *et al*., 2007), were detected in the GeG2^T^ chromid (three GH106 genes) and chromosome (one GH78 and one GH106). The recent isolation and characterization of a α-L-rhamnosidase produced by *Novosphingobium* sp. PP1Y hydrolyzing various glycosylated flavonoids reveals interesting biotechnological potential associated with these enzymes (DeLise *et al*., 2016). Furthermore, CAZymes associated with the biosynthesis of oligosaccharides, polysaccharides and glycoconjugates (GTs) are mainly encoded in the chromosome (Figure 6B). As reported for different EPS producers (Deo *et al*., 2019; Wang *et al*., 2019), most GTs were classified in families GT2 and GT4, which include enzymes with diverse origins and functions (*e. g.* cellulose synthases, chitin synthases, glycosyltransferases) (Breton *et al*., 2006).

Metabolic profiles predicted from KEGG revealed that, as reported for other 22 *Novosphingobium* genomes (Wang et al., 2018), strain GeG2^T^ encodes the complete glycolysis, pentose phosphate and tricarboxylic acid cycle pathways. While many genes involved in carbon metabolism are found both in the chromosome and the chromid (Supplementary Figure S9), genes involved in nitrogen metabolism are encoded exclusively in the chromosome, including extracellular nitrite and nitrate transporter, nitrite reductases (*nirB/nirD*) and nitronate monooxygenase (*ncd2*), as well as genes encoding the Amt family of ammonium transporters.

Interestingly, for some pathways, member genes were found to be interspersed in the chromosome and the chromid. While alkane sulfonate assimilation genes (*ssuABCD*), the only extracellular sulfur assimilation pathway detected in GeG2^T^ genome, were found exclusively in the chromid, genes involved in sulfate, sulfite and sulfide metabolism are encoded solely in the chromosome (Figure 7). Moreover, except for *hisC*, all genes related to histidine biosynthesis (*hisG, hisE, hisJ, hisA, hisF, hisB, hisN, hisD*) are exclusively found in the chromosome, whereas genes encoding enzymes responsible for the conversion of histidine into glutamate (*hutH, hutU, hutI, hutF, hutG*) were detected in the chromid. Similarly, genes involved in *de novo* purine synthesis are encoded in the chromosome and many genes for purine degradation and salvage pathways were identified exclusively in the chromid of strain GeG2^T^. Curiously, even though taurine dioxygenase (*tauD*) was identified in all 27 *Novosphingobium* genomes from diverse habitats analyzed by Kumar et al., 2017, none of the genes associated to the taurine assimilation pathway could be detected in GeG2^T^ genome (Figure 7).

**Figure 7.**
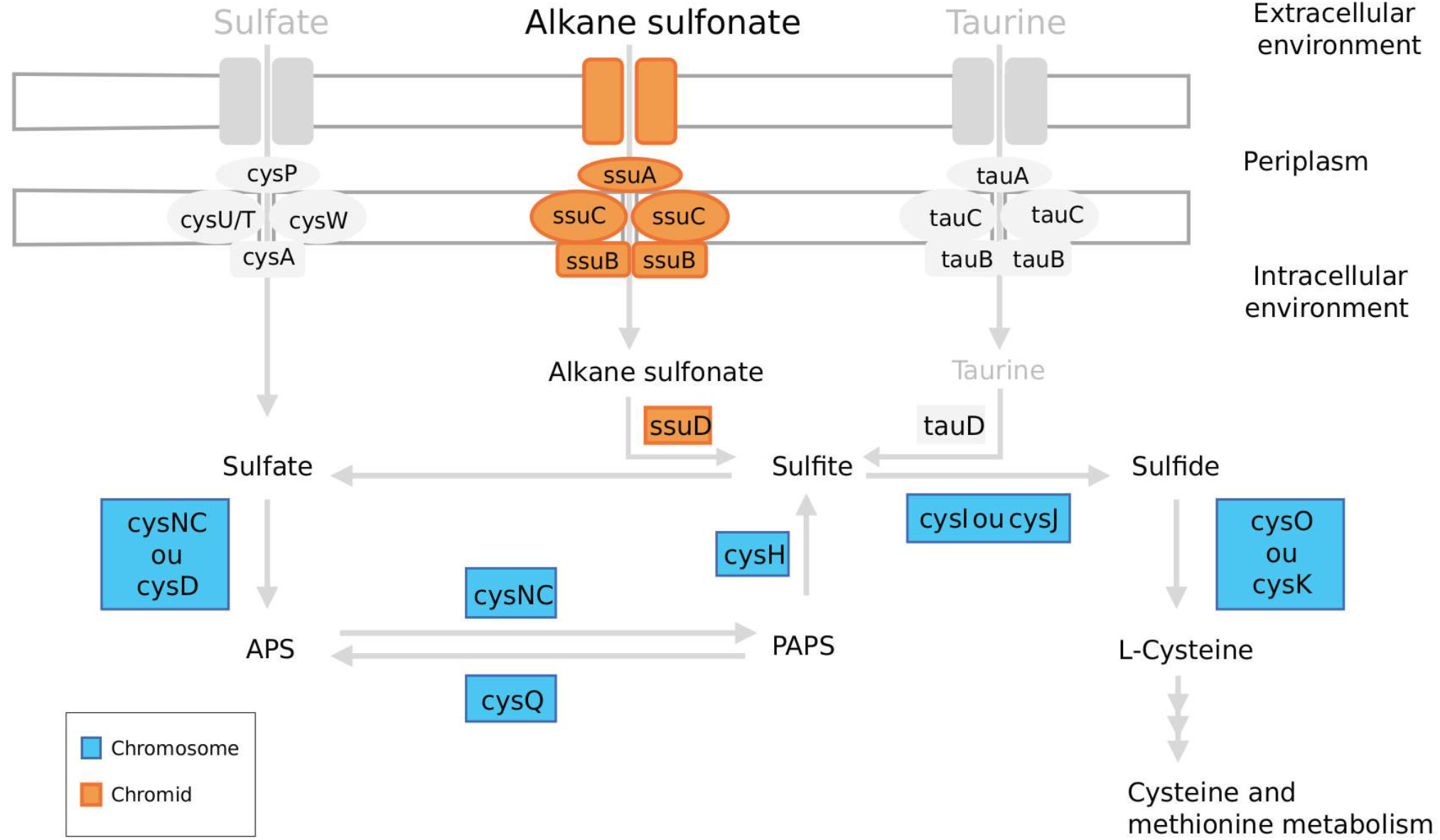
Schematic representation of sulfur metabolism genes identified in strain GeG2^T^ chromosome (in blue) and chromid (in orange). Genes associated with the assimilation of environmental sulfur compounds that were not detected in the genome of strain GeG2^T^ but can be found in other *Novosphingobium* species (Kumar et al., 2017) are represented in grey. APS: adenosine phosphosulfate; PAPS: phosphoadenosine phosphosulfate.

Genes for three secretion systems as well as the twin-arginine translocation (Tat) and secretion (Sec) pathways were detected in the genome of strain GeG2^T^. Interestingly, all genes associated with type IV secretion system (T4SS), Tat and Sec pathways are located in the chromosome, while complete type I and type VI secretion systems (T1SS and T6SS) are coded in the chromid. T4SS have been identified in the chromosome of many bacterial species (Walden et al., 2010; Fischer et al., 2020), including *Sphingomonas* (Wu et al., 2017), and have been associated with horizontal DNA transfer and secretion of bacterial interaction mediators (Alegria *et al*., 2005). T1SS is widely distributed among Gram-negative bacteria and frequently associated with efflux mechanisms that lead to resistance to several compounds in *Sphingomonadaceae* (Wu *et al*., 2017). Although initially identified as a typical virulence factor (Pukatzki *et al*., 2006), T6SS is now recognized as versatile secretion system, commonly found in soil bacteria, with roles in assimilation of scarce ions, inter-bacterial competition, and different interactions with the environment (Coulthurst, 2019). The detection of T1SS and T6SS secretion systems in the chromid of strain GeG2^T^ can further suggest an important role of this replicon in adaptative responses to environmental changes and interactions with other soil microorganisms.

### Aromatic compound degradation

Members of *Novosphingobium* and related genera are recognized for their ability to degrade several xenobiotics and aromatic compounds (Wang *et al*., 2018). A large number of monooxygenases and dioxygenases were detected in strain GeG2^T^ genome (Supplementary Table S2). These enzymes catalyze the ring cleavage step critical to aerobic degradation of aromatic compounds and are essential for the metabolism of a wide variety of recalcitrant substances (Ladino-Orjuela *et al*., 2016). Although additional experimental confirmations are still necessary, taking into consideration the number and classes of monooxygenases and dioxygenases identified in the genome of strain GeG2^T^ (Supplementary Table S2) that are similar to the ones previously reported for *Novosphingobium* species known to be able to degrade compounds such as hexachlorocyclohexane, pentachlorophenol, biphenyl, phenanthrene, pyrene, benzo(a)pyrene, naphthalene, fluorene, among others (Sohn *et al*., 2004; Tiirola *et al*., 2005; Notomista *et al*., 2011; Aylward *et al*., 2013; Saxena *et al*., 2013; Wang *et al*., 2018), strain GeG2^T^ is probably able to metabolize different aromatic compounds.

Interestingly, contrarily to the observed for *N. aromaticivorans* DSM12444, *Novosphingobium* sp. P6W, *Novosphingobium* sp. PP1Y, *N. pentaromativorans* US6-1 and *Novosphingobium* sp. THN1, in which mono and dioxygenases are primarily encoded in chromosomes instead of their secondary megareplicons (Wang *et al*., 2018), most monooxygenases from strain GeG2^T^ are found in the chromid. Dioxygenases, on the other hand, are detected in the chromosome and chromid (Supplementary Table S2). Analysis of known pathways for aromatic compound degradation revealed that, although monooxygenases usually associated with the first steps of toluene and xylene degradation (*tmoA* and *xylM*) could not be annotated in strain GeG2^T^ genome, all other genes for enzymes associated with the biodegradation of these compounds could be identified (Figure 8). Furthermore, many genes from the upper pathway of aromatic compounds degradation producing catechol intermediates were identified both in the chromosome and the chromid, while all genes related to catechol orthoclivage to tricarboxylic acids are encoded in the chromosome (Figure 8).

**Figure 8.**
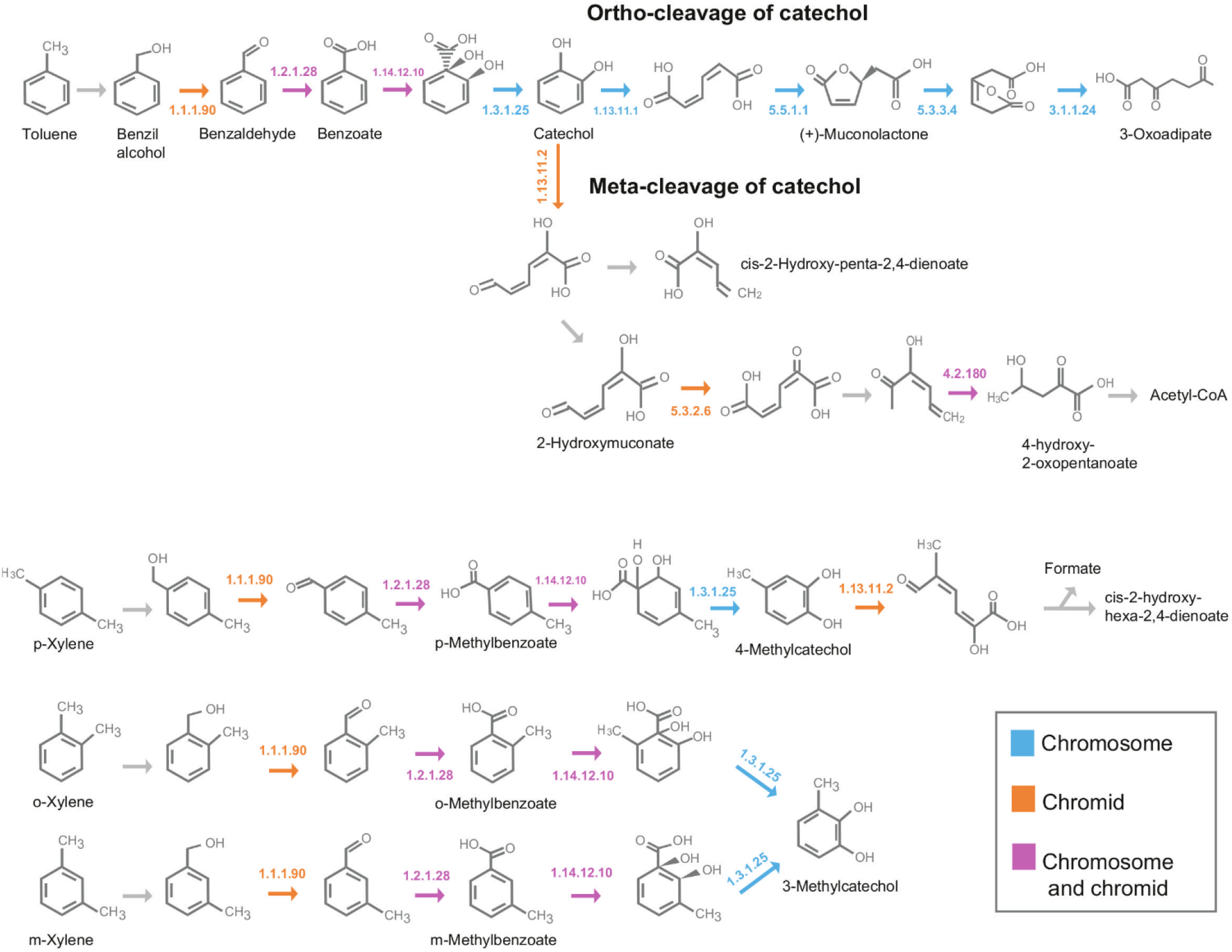
Pathways associated with the degradation of aromatic compounds, obtained with KEGG Mapper tool, indicating genes detected in the chromosome (in blue), the chromid (in orange) or in both replicons (in pink) of strain GeG2^T^ genome. 1.1.1.90: aryl-alcohol dehydrogenase; 1.2.1.28: benzaldehyde dehydrogenase (xylC); 1.14.12.10: benzoate/toluate 1,2-dioxygenase subunit alpha (benA-xylX); 1.3.1.25: dihydroxycyclohexadiene carboxylate dehydrogenase (benD-xylL); 1.13.11.1: catechol 1,2-dioxygenase (catA); 5.5.1.1: muconate cycloisomerase (catB); 5.3.3.4: muconolactone D-isomerase (catC); 3.1.1.24: 3-oxoadipate enol-lactonase (pcaD); 1.13.11.2: catechol 2,3-dioxygenase (dmpB-xylE); 5.3.2.6: 4-oxalocrotonate tautomerase (praC-xylH); 4.2.1.80: 2-keto-4-pentenoate hydratase (mhpD).

## CONCLUSION

In this study, we isolated a novel bacterial strain (GeG2^T^) from soils of a native area of Cerrado, a highly biodiverse biome located in Central Brazil. Based on 16S rRNA gene sequence, strain GeG2^T^ belongs to the alphaproteobacterial genus *Novosphingobium*, presenting 100% nucleotide identity (95% coverage) with previously characterized species *Novosphingobium rosa*. Albeit this fact, we conducted thorough morphological, biochemical and genomic analyses to describe GeG2^T^. This strain presented planktonic-sessile dimorphism and microscopical analyses indicated the production of exopolysaccharide, variable sized extracellular vesicles as well as intracytoplasmic electron-dense granules with characteristic aspect of polyphosphate granules or acidocalcisomes. The complete genome of GeG2^T^ was resolved through a combination of long and short-reads sequencing approaches revealing a multipartite architecture consisting of a chromosome (4.2 Mb), an extrachromosomal megareplicon (2.7 Mb) and two plasmids (212 and 68 kb). Although experimental demonstrations would be ideally necessary for a definitive classification of the secondary megareplicon identified in GeG2^T^, the identification of chromosomal signatures and plasmid-type replication and maintenance systems indicates that it is a chromid (diCenzo and Finan, 2017). Genome-based taxonomic identification, including overall genome relatedness index (OGRI) estimations and phylogenomic analysis, indicated a clear distinction between strain GeG2^T^ and other *Novosphingobium* representatives, even *N. rosa*, despite their 100% rRNA gene similarity. Furthermore, whole genome sequence alignment between strain GeG2^T^ and *N. rosa* NBRC15208 further demonstrated their dissimilarity. In terms of gene space, a broad spectrum of carbohydrate metabolism enzymes and the large number of monooxygenases and dioxygenases identified in the genome of strain GeG2^T^ reveal its great potential for plant biomass degradation, polysaccharide production and degradation of diverse aromatic compounds. In summary, a polyphasic characterization, including physiologic, chemotaxonomic, MALFI-TOF protein profile and whole genome-based analyses, indicates that strain GeG2^T^ represents a new species within *Novosphingobium* genus for which the name *Novosphingobium terrae* sp. nov. is proposed.

### Description of *Novosphingobium terrae* sp. nov

*Novosphingobium terrae* (ter′rae. L. gen. n. *terrae* of soil). Cells are Gram-negative bacilli, with 0.3 - 0.6 µm in width and 1.3-2.3 µm in length. Colonies grown in minimal media (MM) were white, circular, convex, with regular edges, shiny appearance and about 2-3 mm diameter. Growth occurs at 15–33 °C, pH 4-7 and NaCl concentrations from 0 to 1% (w/v). Positive for aesculin hydrolysis, catalase and oxidase activity, assimilation of glucose, arabinose, mannose, N-acetyl-glucosamine, maltose, gluconate and activities of alkaline phosphatase, leucin-arylamidase, valin-arylamidase, acid phosphatase, naphtol-AS-BI-phosphohydrolase, β-galactosidase, α-glucosidase and β-glucosidase. Weakly positive for nitrate reduction. Negative for hydrolysis of gelatin and urea; indol production; glucose fermentation; assimilation of mannitol, caprate, adipate, malate, citrate, phenylacetate; arginine dihydrolase, lipase (C14) and α-mannosidase activities.

The type strain, GeG2^T^ (=CBMAI 2313^T^ =CBAS 753^T^) was isolated from Cerrado soils collected in Brasilia, Brazil (15°55’ S, 47°51’ W). The DNA GC content of the type strain is 63.57 mol %. The 16S rRNA gene sequence (MT325926.1) and the complete genome sequence of *Novosphingobium terrae* GeG2^T^ (GCA_017163935.1) have been deposited in GenBank.

## ACKNOWLEDGMENTS

The authors would like to thank Dr. Heloisa Sinatora Miranda for assistance in soil collections. We acknowledge the Center of Microscopy at Federal University of Minas Gerais (http://www.microscopia.ufmg.br) and the Laboratory of Microscopy at University of Brasilia for providing the equipment and technical support for experiments involving transmission electron microscopy and scanning electron microscopy, respectively. We would also like to thank the Laboratory of Mass Spectrometry at Embrapa for providing the equipment and support for MALDI Biotyper analysis.

## AUTHOR CONTRIBUTIONS

AB cultivated and isolated the bacterial strain. AB and CV performed the microbiological characterizations. FA, RR, AB, RK and GP analyzed the sequence data. AB and MR performed MALDI-TOF analyses. MT performed fatty acid analysis. CK and RK acquired funding for this study. AB, FA, CK and GP wrote the manuscript. All authors revised and approved the manuscript.

## DATA AVAILABILITY STATEMENT

The whole genome sequencing dataset is available under the NCBI Bioproject PRJNA624997.

## REFERENCES

Agustini, B.C., Silva, L.P., Bloch, Jr C., Bonfim, T.M., da Silva, G.A. (2014). Evaluation of MALDI-TOF mass spectrometry for identification of environmental yeasts and development of supplementary Database. Appl. Microbiol. Biotechnol., 98:5645–5654. doi: 10.1007/s00253-014-5686-7

Alegria, M.C., Souza, D.P., Andrade, M.O. Docena, C., Khater, L., Ramos, C.H., et al. (2005). Identification of new protein-protein interactions involving the products of the chromosome- and plasmid-encoded type IV secretion loci of the phytopathogen *Xanthomonas axonopodis* pv. citri. J. Bacteriol. 187, 2315–2325. doi: 10.1128/JB.187.7.2315-2325.2005

Alves-Prado, H.F.; Pavezzi, F.C.; Leite, R.S. de Oliveira, V.M., Sette, L.D., Dasilva, R. (2010). Screening and production study of microbial xylanase producers from Brazilian Cerrado. Appl. Biochem. Biotechnol., 161(1-8):333–46. doi: 10.1007/s12010-009-8823-5

Andrews, S. (2010). FastQC: A Quality Control Tool for High Throughput Sequence Data. Available online at: http://www.bioinformatics.babraham.ac.uk/projects/fastqc/

Aramaki, T., Blanc-Mathieu, R., Endo, H., Ohkubo, K., Kanehisa, M., Goto, S., Ogata, H. (2020). KofamKOALA: KEGG Ortholog assignment based on profile HMM and adaptive score threshold. Bioinformatics, 36(7):2251–2252. doi: 10.1093/bioinformatics/btz859

Aylward, F.O., McDonald, B.R., Adams, S.M., Valenzuela, A., Schmidt, R.A., Goodwin, L.A., et al. (2013). Comparison of 26 sphingomonad genomes reveals diverse environmental adaptations and biodegradative capabilities. Appl. Environ. Microbiol.,79(12):3724–3733. doi: 10.1128/AEM.00518-13

Baek, S.H., Lim, J.H., Jin, L., Lee, H.G., Lee, S.T. (2011). *Novosphingobium sediminicola* sp. nov. isolated from freshwater sediment. Int. J. Syst. Evol. Microbiol., 61(Pt 10):2464–2468. doi: 10.1099/ijs.0.024307-0

Bauer, A.W., Kirby, W.M.M., Sherris, J.C., Turck, M. (1966). Antibiotic Susceptibility Testing by a Standardized Single Disk Method. Am. J. Clin. Pathol., 45(4):493–496. doi: 10.1093/ajcp/45.4_ts.493

Breton, C., Snajdrová, L., Jeanneau, C., Koca, J., Imberty, A. (2006). Structures and mechanisms of glycosyltransferases. Glycobiology, 16(2):29R–37R. doi: 10.1093/glycob/cwj016

Busse, H.J., Kämpfer, P., Denner, E.B. (1999). Chemotaxonomic characterisation of *Sphingomonas*. J. Ind. Microbiol. Biotechnol., 23(4-5):242–251. doi: 10.1038/sj.jim.2900745

Carver, T., Thomson, N., Bleasby, A., Berriman, M., Parkhill, J. (2009). DNAPlotter: circular and linear interactive genome visualization. Bioinformatics, 25(1):119–120 doi: 10.1093/bioinformatics/btn578

Chain, P.S., Denef, V.J., Konstantinidis, K.T., Vergez, L.M., Agulló, L., Reyes, V.L. (2006). *Burkholderia xenovorans* LB400 harbors a multi-replicon, 9.73-Mbp genome shaped for versatility. Proc. Natl. Acad. Sci. USA., 103(42):15280–15287. doi: 10.1073/pnas.0606924103

Choi, C.W., Park, E.C., Yun, S.H., Lee, S.Y., Lee, Y.G., Hong, Y. et al. (2014). Proteomic characterization of the outer membrane vesicle of *Pseudomonas putida* KT2440. J. Proteome Res., 13(10):4298–4309. doi: 10.1021/pr500411d

Chun, J., Oren, A., Ventosa, A., Christensen, H., Arahal, D.R., da Costa, M.S. et al. (2018). Proposed minimal standards for the use of genome data for the taxonomy of prokaryotes. Int. J. Syst. Evol. Microbiol., 68(1):461–466. doi: 10.1099/ijsem.0.002516

Coppotelli, B.M., Ibarrolaza, A., Dias, R.L., Del Panno, M.T., Berthe-Corti, L., Morelli, I.S. (2010). Study of the degradation activity and the strategies to promote the bioavailability of phenanthrene by *Sphingomonas paucimobilis* strain 20006FA. Microb. Ecol., 59(2):266–276. doi: 10.1007/s00248-009-9563-3

Coulthurst, S. (2019). The Type VI secretion system: a versatile bacterial weapon. Microbiology, 165(5):503–515. doi: 10.1099/mic.0.000789

Dantas, G., Sommer, M.O., Oluwasegun, R.D., Church, G.M. (2008). Bacteria subsisting on antibiotics. Science, 320(5872):100–103. doi: 10.1126/science.1155157

D’Argenio, V., Notomista, E., Petrillo, M., Cantiello, P., Cafaro, V., Izzo, V. et al. (2014). Complete sequencing of *Novosphingobium* sp. PP1Y reveals a biotechnologically meaningful metabolic pattern. BMC Genomics, 15(1):384. doi: 10.1186/1471-2164-15-384

De Lise, F., Mensitieri, F., Rusciano, G., Dal Piaz, F., Forte, G., Di Lorenzo, F. et al. (2019). *Novosphingobium* sp. PP1Y as a novel source of outer membrane vesicles. J Microbiol., 57(6):498–508. doi: 10.1007/s12275-019-8483-2

De Lise, F., Mensitieri, F., Tarallo, V., Ventimiglia, N., Vinciguerra, R., Tramice, A. et al. (2016). RHA-P. Isolation, expression and characterization of a bacterial α-l-rhamnosidase from *Novosphingobium* sp. PP1Y. J. Mol. Catalysis B: Enzymatic, 134:136–147. doi: 10.1016/j.molcatb.2016.10.002

Deo, D., Davray, D., Kulkarni, R. (2019). A Diverse Repertoire of Exopolysaccharide Biosynthesis Gene Clusters in Lactobacillus Revealed by Comparative Analysis in 106 Sequenced Genomes. Microorganisms, 7(10), 444. doi: 10.3390/microorganisms7100444

DiCenzo, G.C., and Finan, T.M. (2017). The Divided Bacterial Genome: Structure, Function, and Evolution. Microbiol. Mol. Biol. Rev., 81(3):e00019–17. doi: 10.1128/MMBR.00019-17

Docampo, R. (2006). Acidocalcisomes and Polyphosphate Granules. In: Inclusions in Prokaryotes - Microbiology Monographs, vol 1. Springer, Berlin, Heidelberg. doi: 10.1007/3-540-33774-1_3

Fischer, W., Tegtmeyer, N., Stingl, K., Backert, S. (2020). Four Chromosomal Type IV Secretion Systems in Helicobacter pylori: Composition, Structure and Function. Front. Microbiol., 11, 1592. doi: 10.3389/fmicb.2020.01592

Fox, G.E., Wisotzkey, J.D., Jurtshuk Jr, P. (1992) How close is close: 16S rRNA sequence identity may not be sufficient to guarantee species identity. Int. J. Syst. Bacteriol., 42(1):166–170. doi: 10.1099/00207713-42-1-166

Frank, C., Jendrossek, D. (2020). Acidocalcisomes and Polyphosphate Granules Are Different Subcellular Structures in Agrobacterium tumefaciens. Appl. Environ. Microbiol., 86(8):e02759–19. doi: 10.1128/AEM.02759-19

Frank, O., Göker, M., Pradella, S., Petersen, J. (2015). Ocean’s twelve: flagellar and biofilm chromids in the multipartite genome of *Marinovum algicola* DG898 exemplify functional compartmentalization. Environ. Microbiol., 17:4019–4034. doi: 10.1111/1462-2920.12947

Gilewicz, M., Ni’matuzahroh, Nadalig, T., Budzinski, H., Doumenq, P., Michotey, V., et al. (1997). Isolation and characterization of a marine bacterium capable of utilizing 2-methylphenanthrene. Appl. Microbiol. Biotechnol., 48, 528–533. doi: 10.1007/s002530051091

Glaeser, S.P., Bolte, K., Martin, K., Busse, H.J., Grossart, H.P., Kämpfer, P, Glaeser J. (2013a). *Novosphingobium fuchskuhlense* sp. nov., isolated from the north-east basin of Lake Grosse Fuchskuhle. Int. J. Syst. Evol. Microbiol.,, 63(Pt 2):586–592. doi: 10.1099/ijs.0.043083-0

Gogoleva, N.E., Nikolaichik, Y.A., Ismailov, T.T. Gorshkov, V.Y., Safronova, V.I., Belimov, A.A., Gogolev, Y. (2019). Complete genome sequence of the abscisic acid-utilizing strain *Novosphingobium* sp. P6W. 3 Biotech., 9(3):94. doi: 10.1007/s13205-019-1625-8

Glaeser, S.P., and Kämpfer, P. (2014) The Family *Sphingomonadaceae*. *In:* The Prokaryotes. Springer, Berlin, Heidelberg. doi: 10.1007/978-3-642-30197-1_302

Goutx M., Mutaftshiev, S., Bertrand, J. (1987). Lipid and exopolysaccharide production during hydrocarbon growth of a marine bacterium from the sea surface. Mar. Ecol. Prog. Ser., 40(3), 259–265. doi: 10.3354/meps040259

Gupta, S.K., Lal, D., Lal, R. (2009). *Novosphingobium panipatense* sp. nov. and *Novosphingobium mathurense* sp. nov., from oil-contaminated soil. Intern. J. Syst. Bacteriol., 59, 156–161. doi: 10.1099/ijs.0.65743-0

Gurevich, A., Saveliev, V., Vyahhi, N., Tesler, G. (2013). QUAST: quality assessment tool for genome assemblies. Bioinformatics, 29(8):1072–1075. doi: 10.1093/bioinformatics/btt086

Ha, S.M., Kim, C.K., Roh, J., Byun, J.H., Yang, S.J., Choi, S.B. et al. (2019). Application of the Whole Genome-Based Bacterial Identification System, TrueBac ID, Using Clinical Isolates That Were Not Identified With Three Matrix-Assisted Laser Desorption/Ionization Time-of-Flight Mass Spectrometry (MALDI-TOF MS) Systems. Ann. Lab. Med., 39(6), 530–536. doi: 10.3343/alm.2019.39.6.530

Haridasan M. (2008). Nutritional adaptations of native plants of the cerrado biome in acid soils. *Braz*. J. Plant Physiol, 20(3), 183–195. doi: 10.1590/S1677-04202008000300003

Harrison, P.W., Lower, R.P., Kim, N.K., Young, J.P. (2010). Introducing the bacterial ’chromid’: not a chromosome, not a plasmid. Trends Microbiol., 18(4):141–148. doi: 10.1016/j.tim.2009.12.010

Hegedűs, B., Kós, P.B., Bálint, B., Maróti, G., Gan, H.M. et al (2017) Complete genome sequence of *Novosphingobium resinovorum* SA1, a versatile xenobiotic-degrading bacterium capable of utilizing sulfanilic acid. J Biotechnol., 241:76–80. doi: 10.1016/j.jbiotec.2016.11.013

Huerta-Cepas, J., Forslund, K., Coelho, L.P., Szklarczyk, D., Jensen, L.J., von Mering, C., Bork, P. (2017). Fast Genome-Wide Functional Annotation through Orthology Assignment by eggNOG- Mapper. Mol. Biol. Evol. 34(8):2115–2122. doi: 10.1093/molbev/msx148

Huo, Y.Y., You, H., Li, Z.Y., Wang, C.S., Xu, X.W. (2015). *Novosphingobium marinum* sp. nov., isolated from seawater. Int. J. Syst. Evol. Microbiol.,65(Pt 2):676–80. doi: 10.1099/ijs.0.070433-0

Husain, D.R., Goutx, M., Bezac, C., Gilewicz, M. and Bertrand, J.-C. (1997), Morphological adaptation of *Pseudomonas nautica* strain 617 to growth on eicosane and modes of eicosane uptake. Letters Appl. Microbiol., 24: 55–58. doi: 10.1046/j.1472-765X.1997.00345.x

Jain, C., Rodriguez-R, L.M., Phillippy, A.M., Konstantinidis, K.T., Aluru, S. (2018). High throughput ANI analysis of 90K prokaryotic genomes reveals clear species boundaries. Nat Commun., 9(1):5114. doi: 10.1038/s41467-018-07641-9

Janssen, P.J., Van Houdt, R., Moors, H., Monsieurs, P., Morin, N., Michaux, A. et al. (2010). The complete genome sequence of *Cupriavidus metallidurans* strain CH34, a master survivalist in harsh and anthropogenic environments. PLoS One, 5(5):e10433. doi: 10.1371/journal.pone.0010433

Jaspers, E., and Overmann, J. (2004). Ecological significance of microdiversity: identical 16S rRNA gene sequences can be found in bacteria with highly divergent genomes and ecophysiologies. Appl. Environ. Microbiol., 70(8):4831–4839. doi: 10.1128/AEM.70.8.4831-4839.2004

Kämpfer, P., Busse, H.J., Glaeser, S.P. (2018). *Novosphingobium lubricantis* sp. nov., isolated from a coolant lubricant emulsion. Int. J. Syst. Evol. Microbiol., 68(5):1560–1564. doi: 10.1099/ijsem.0.002702

Kämpfer, P., Denner, E.B., Meyer, S., Moore, E.R., Busse, H.J. (1997) Classification of “*Pseudomonas azotocolligans*” in the genus *Sphingomonas* as *Sphingomonas trueperi* sp. nov. Int. J. Syst. Bacteriol. 47:577–583. doi: 10.1099/00207713-47-2-577

Kämpfer, P., Young, C.C., Busse, H., Lin, S.Y., Rekha, P.D., Arun, A.B. et al. (2011). *Novosphingobium soli* sp. nov., isolated from soil. Int J Syst Evol Microbiol., 61(Pt 2):259–263. doi: 10.1099/ijs.0.022178-0

Kämpfer, P., Martin, K., McInroy, J.A., Glaeser, S.P. (2015). Proposal of *Novosphingobium rhizosphaerae* sp. nov., isolated from the rhizosphere. Int. J. Syst. Evol. Microbiol., 65(Pt 1):195–200. doi: 10.1099/ijs.0.070375-0

Kanehisa, M., Araki, M., Goto, S., Hattori, M., Hirakawa, M., Itoh, M. et al. (2008). KEGG for linking genomes to life and the environment. Nucleic Acids Res., 36:D480–D484. doi: 10.1093/nar/gkm882

Kimura, M. (1980). A simple method for estimating evolutionary rates of base substitutions through comparative studies of nucleotide sequences. J. Mol. Evol. 16(2):111–120. doi: 10.1007/BF01731581

Krawczyk, P.S., Lipinski, L., Dziembowski, A. (2018). PlasFlow: predicting plasmid sequences in metagenomic data using genome signatures. Nucleic Acids Res., 46(6):e35. doi: 10.1093/nar/gkx1321

Kumar, R., Verma, H., Heider, S., Bajaj, A., Sood, U., Ponnusamy, K. et al. (2017). Comparative genomic analysis reveals habitat-specific genes and regulatory hubs within the genus *Novosphingobium*. mSystems 2, e00020–17, doi: 10.1128/mSystems.00020-17

Kurtz, S., Phillippy, A., Delcher, A.L., Smoot, M., Shumway, M., Antonescu, C., Salzberg, S.L. (2004). Versatile and open software for comparing large genomes. Genome Biol., 5(2):R12. doi: 10.1186/gb-2004-5-2-r12

Ladino-Orjuela, G., Gomes, E., da Silva, R., Salt, C., Parsons, J.R. (2016) Metabolic Pathways for Degradation of Aromatic Hydrocarbons by Bacteria. In: Reviews of Environmental Contamination and Toxicology Volume 237, Springer, Cham. doi: 10.1007/978-3-319-23573-8_5

Lane, D.J. (1991). 16S/23S rRNA sequencing in Nucleic acid techniques in bacterial systematics. New York: John Wiley and Sons. P. 115–175.

Lee, J.C., Kim, S.G., Whang, K.S. (2014a). *Novosphingobium aquiterrae* sp. nov., isolated from ground water. Int J Syst Evol Microbiol.,64(Pt 9):3282–3287. doi: 10.1099/ijs.0.060749-0

Lee, L.H., Azman, A.S., Zainal, N., Eng, S.K., Fang, C.M., Hong, K., Chan, K.G. (2014b). *Novosphingobium malaysiense* sp. nov. isolated from mangrove sediment. Int. J Syst. Evol. Microbiol. 64(pt4):1194–1201. doi: 10.1099/ijs.0.059014-0

Li, X.Z., and Nikaido, H. (2009). Efflux-mediated drug resistance in bacteria: an update. Drugs, 69(12), 1555–1623. doi: 10.2165/11317030-000000000-00000

Mackenzie, C., Choudhary, M., Larimer, F.W., Predki, P.F., Stilwagen, S., Armitage, J.P. et al. (2001). The home stretch, a first analysis of the nearly completed genome of Rhodobacter sphaeroides 2.4.1. Photosynth. Res., 70(1):19–41. doi: 10.1023/A:1013831823701

Manzanares, P., Vallés, S., Ramòn, D., Orejas, M. (2007) α-L-rhamnosidases: Old and New Insights. *In*: Industrial Enzymes. Springer, Dordrecht. doi: 10.1007/1-4020-5377-0_8

Meier-Kolthoff, J.P., and Göker, M. (2019). TYGS is an automated high-throughput platform for state-of-the-art genome-based taxonomy. Nat. Commun., 10:2182. doi: 10.1038/s41467-019-10210-3

Ngo, H.T., Trinh, H., Kim, J.H., Yang, J.E., Won, K.H., Kim, J.H. et al. (2016). *Novosphingobium lotistagni* sp. nov., isolated from a lotus pond. Int. J. Syst. Evol. Microbiol., 66(11):4729–4734. doi: 10.1099/ijsem.0.001418

Nguyen, S.T.C., Freund, H.L., Kasanjian, J., Berlemont, R. (2018). Function, distribution, and annotation of characterized cellulases, xylanases, and chitinases from CAZy. Appl. Microbiol. Biotechnol., 102(4):1629–1637. doi: 10.1007/s00253-018-8778-y

Notomista, E., Pennacchio, F., Cafaro, V., Smaldone, G., Izzo, V., Troncone, L. et al. (2011). The marine isolate *Novosphingobium* sp. PP1Y shows specific adaptation to use the aromatic fraction of fuels as the sole carbon and energy source. Microb. Ecol., 61(3):582–594. doi: 10.1007/s00248-010-9786-3

Parks, D.H., Imelfort, M., Skennerton, C.T., Hugenholtz, P., Tyson, G.W. (2014). CheckM: Assessing the quality of microbial genomes recovered from isolates, single cells, and metagenomes. Genome Res., 25: 1043–1055. doi: 10.1101/gr.186072.114

Peixoto, J., Silva, L.P., Krüger, R. H. (2017). Brazilian Cerrado soil reveals an untapped microbial potential for unpretreated polyethylene biodegradation. J. Hazard. Mater., 15,324(Pt B):634–644. doi: 10.1016/j.jhazmat.2016.11.037

Pollock, T., and Armentrout, R. (1999). Planktonic/sessile dimorphism of polysaccharide-encapsulated sphingomonads. J. Ind. Microbiol. Biotech., 23, 436–441. doi: 10.1038/sj.jim.2900710

Pukatzki, S., Ma, A.T., Sturtevant, D., Krastins, B., Sarracino, D., Nelson, W.C. et al. (2006) Identification of a conserved bacterial protein secretion system in Vibrio cholerae using the Dictyostelium host model system. Proc. Natl. Acad. Sci. USA, 103(5):1528–1533. doi: 10.1073/pnas.0510322103

Raina, V., Nayak, T., Ray, L., Kumari, K., Suar, M. (2019). Approach for designation and description of novel microbial species. In: Microbial diversity in the genomic era (pp. 137–152). India: Elsevier.

Ramasamy, D., Mishra, A.K., Lagier, J.C., Padhmanabhan, R., Rossi M., Sentausa, E. et al. (2014). A polyphasic strategy incorporating genomic data for the taxonomic description of novel bacterial species. Int. J. Syst. Evol. Microbiol. 64, 384–391. 10.1099/ijs.0.057091-0

Sasser, M. (1990). Identification of bacteria by gas chromatography of cellular fatty acids. MIDI Technical Note (MIDI, Newark, Delaware),101, 1–7.

Saxena, A., Anand, S., Dua, A., Sangwan, N., Khan, F., Lal, R. (2013). *Novosphingobium lindaniclasticum* sp. nov., a hexachlorocyclohexane (HCH)-degrading bacterium isolated from an HCH dumpsite. Int. J. Syst. Evol. Microbiol.,63(Pt 6):2160–2167. doi: 10.1099/ijs.0.045443-0

Schwechheimer, C., and Kuehn, M.J. (2015). Outer-membrane vesicles from Gram-negative bacteria: biogenesis and functions. Nat. Rev. Microbiol., 13(10):605–619. doi: 10.1038/nrmicro3525

Seemann, T. (2014) Prokka: rapid prokaryotic genome annotation. Bioinformatics, 30(14):2068–9. doi: 10.1093/bioinformatics/btu153.

Sha, S., Zhong, J., Chen, B., Lin, L., Luan, T. (2017). *Novosphingobium guangzhouense* sp. nov., with the ability to degrade 1-methylphenanthrene. Int. J. Syst. Evol. Microbiol., 67(2):489–497. doi: 10.1099/ijsem.0.001669

Sheu, S.Y., Cai, C.Y., Kwon, S.W., Chen, W.M. (2020). *Novosphingobium umbonatum* sp. nov., isolated from a freshwater mesocosm. Int. J. Syst. Evol. Microbiol., 70(2):1122-1132. doi: 10.1099/ijsem.0.003889

Sheu, S.Y., Chen, Z.H., Chen, W.M. (2016). *Novosphingobium piscinae* sp. nov., isolated from a fish culture pond. Int. J. Syst. Evol. Microbiol., 66(3):1539–1545. doi: 10.1099/ijsem.0.000914

Sheu, S.Y., Huang, C.W., Chen, J.C., Chen, Z.H., Chen, W M. (2018). *Novosphingobium arvoryzae* sp. nov., isolated from a flooded rice field. Int. J Syst Evol Microbiol. 68, 2151–2157. doi: 10.1099/ijsem.0.002756

Sohn, J.H., Kwon, K.K., Kang, J.H., Jung, H.B., Kim, S.J. (2004). *Novosphingobium pentaromativorans* sp. nov., a high-molecular-mass polycyclic aromatic hydrocarbon-degrading bacterium isolated from estuarine sediment. Int. J. Syst. Evol. Microbiol., 54(Pt 5):1483–1487. doi: 10.1099/ijs.0.02945-0.

Souza, W. (2007). Técnicas de microscopia eletrônica aplicadas às ciências biológicas. 3a edição. Sociedade Brasileira de Microscopia, Rio de Janeiro.

Stackebrandt, E., and Goebel, BM. (1994). Taxonomic Note: A Place for DNA-DNA Reassociation and 16S rRNA Sequence Analysis in the Present Species Definition in Bacteriology. Int. J. Syst. Evol. Microbiol., 44(4):846–849. doi: 10.1099/00207713-44-4-846

Suzuki, Y., Nishijima, S., Furuta, Y., Yoshimura, J., Suda, W., Oshima, K. et al. (2019). Long-read metagenomic exploration of extrachromosomal mobile genetic elements in the human gut. Microbiome, 7(1):119. doi: 10.1186/s40168-019-0737-z

Takeuchi, M., Hamana, K., Hiraishi, A. (2001). Proposal of the genus *Sphingomonas sensu stricto* and three new genera, *Sphingobium, Novosphingobium* and *Sphingopyxis*, on the basis of phylogenetic and chemotaxonomic analyses. Intern. J. System. Bacteriol., 51(Pt 4):1405–1417. doi: 10.1099/00207713-51-4-1405

Takeuchi, M., Sakane, T., Yanagi, M., Yamasato, K., Hamana, K., Yokota, A. (1995). Taxonomic Study of Bacteria Isolated from Plants: Proposal of *Sphingomonas rosa* sp. nov., *Sphingomonas pruni* sp. nov., *Sphingomonas asaccharolytica* sp. nov., and *Sphingomonas mali* sp. nov. Int J Syst Bacteriol., 45(2):334–341. doi: 10.1099/00207713-45-2-334

Tamura, K., Peterson, D., Peterson, N., Stecher, G., Nei, M., Kumar, S. (2011). MEGA5: Molecular evolutionary genetics analyses using maximum likelihood, evolutionary distance and maximum parsimony methods. Mol. Biol. Evol., 28: 2731–2739. doi: 10.1093/molbev/msr121

Tamura, K., Nei, M., Kumar, S. (2004). Prospects for inferring very large phylogenies by using the neighbor-joining method. Proc. Natl. Acad. Sci. USA, 101:11030–11035.

Tiirola, M.A., Busse, H.J., Kämpfer, P., Männistö, M.K. (2005). *Novosphingobium lentum* sp. nov., a psychrotolerant bacterium from a polychlorophenol bioremediation process. Int J Syst Evol Microbiol. 55(Pt 2):583–588. doi: 10.1099/ijs.0.63386-0

Tiirola, M.A., Männistö, M.K., Puhakka, J.A., Kulomaa, M.S. (2002). Isolation and characterization of *Novosphingobium* sp. strain MT1, a dominant polychlorophenol-degrading strain in a groundwater bioremediation system. Appl. Environ. Microbiol., 68(1):173–180. doi: 10.1128/aem.68.1.173-180.2002

Tindall, B.J. (1990a). A comparative study of the lipid composition of *Halobacterium saccharovorum* from various sources. Syst. Appl. Microbiol,. 13, 128–130. doi: 10.1016/S0723-2020(11)80158-X

Tindall, B.J. (1990b). Lipid composition of *Halobacterium lacusprofundi*. FEMS Microbiol. Letts., 66, 199–202. doi: 10.1016/0378-1097(90)90282-U

Tindall, B.J., Sikorski, J., Smibert, R.A., Krieg, N.R. (2007), Phenotypic characterization and the principles of comparative systematics. In: Methods for General and Molecular Bacteriology, 3rd edn. pp330–393. Washington, D.C. American Society for Microbiology.

Toyofuku, M., Nomura, N., Eberl, L. (2019). Types and origins of bacterial membrane vesicles. Nat. Rev. Microbiol., 17, 13–24.doi: 10.1038/s41579-018-0112-2

Wallden, K., Rivera-Calzada, A., Waksman, G. (2010). Type IV secretion systems: versatility and diversity in function. Cell. Microbiol., 12(9), 1203–1212. doi: 10.1111/j.1462-5822.2010.01499.x

Wang, J., Wang, C., Li, J., Bai, P., Li, Q., Shen, M. et al. (2018). Comparative Genomics of Degradative *Novosphingobium* Strains With Special Reference to Microcystin-Degrading *Novosphingobium* sp. THN1. Front. Microbiol., 9:2238. doi: 10.3389/fmicb.2018.02238

Wang, J., Salem, D.R., Sani, R.K. (2019). Extremophilic exopolysaccharides: A review and new perspectives on engineering strategies and applications. Carbohydr Polym., 205:8–26. doi: 10.1016/j.carbpol.2018.10.011

Waterhouse, R.M., Seppey, M., Simão, F.A., Manni, M., Ioannidis, P., Klioutchnikov, G., et al. (2018). BUSCO Applications from Quality Assessments to Gene Prediction and Phylogenomics. Mol Biol Evol., 35(3):543–548. doi: 10.1093/molbev/msx319

Wick, R.R., Judd, L.M., Gorrie, C.L., Holt, K.E. (2017) Unicycler: Resolving bacterial genome assemblies from short and long sequencing reads. PloS Comput. Biol., 13(6): e1005595. doi: 10.1371/journal.pcbi.1005595

Wu, M., Huang, H., Li, G., Ren, Y., Shi, Z., Xiaoyan, L. et al. (2017). The evolutionary life cycle of the polysaccharide biosynthetic gene cluster based on the Sphingomonadaceae. Sci. Rep., 7, 46484. doi: 10.1038/srep46484

Yabuuchi, E., Kosako, Y., Fujiwara, N., Naka, T., Matsunaga, I., Ogura, H., Kobayashi, K. (2002) Emendation of the genus Sphingomonas Yabuuchi et al. 1990 and junior objective synonymy of the species of three genera, *Sphingobium*, *Novosphingobium* and *Sphingopyxis*, in conjunction with *Blastomonas ursincola*. Int. J. Syst. Evol. Microbiol., 52(Pt 5):1485–1496. doi: 10.1099/00207713-52-5-1485

Yin, T., Cook, D., Lawrence, M. (2012). ggbio: an R package for extending the grammar of graphics for genomic data. Genome Biol., 13(8):R77. doi: 10.1186/gb-2012-13-8-r77

Yoon, S.H., Ha, S.M., Kwon, S., Lim, J., Kim, Y., Seo, H., Chun, J. (2017). Introducing EzBioCloud: A taxonomically united database of 16S rRNA and whole genome assemblies. Int. J. Syst. Evol. Microbiol., 67:1613–1617. doi: 10.1099/ijsem.0.001755

Yuan, J., Lai, Q., Zheng, T., Shao, Z. (2009). *Novosphingobium indicum* sp. nov., a polycyclic aromatic hydrocarbon-degrading bacterium isolated from a deep-sea environment. Int J Syst Evol Microbiol., 59(Pt 8):2084–2088. doi: 10.1099/ijs.0.002873-0

Yun, S.H., Lee, S.Y., Choi, C.W., Lee, H., Ro, H.J., Jun, S., et al. (2017). Proteomic characterization of the outer membrane vesicle of the halophilic marine bacterium *Novosphingobium pentaromativorans* US6-1. J. Microbiol., 55(1):56–62. doi: 10.1007/s12275-017-6581-6

Zhang, J., Kobert, K., Flouri, T., Stamatakis, A. (2014). PEAR: a fast and accurate Illumina Paired- End reAd mergeR. Bioinformatics, 30(5), 614–620. doi: 10.1093/bioinformatics/btt593

Zhang, H., Yohe, T., Huang, L., Entwistle, S., Wu, P., Yang, Z. et al. (2018). dbCAN2: a meta server for automated carbohydrate-active enzyme annotation. Nucleic Acids Res., 46(W1):W95–W101. doi: 10.1093/nar/gky418

Zheng, J., Guan, Z., Cao, S., Peng, D., Ruan, L., Jiang, D., Sun, M. (2015). Plasmids are vectors for redundant chromosomal genes in the *Bacillus cereus* group. BMC Genomics, 16(1):6. doi: 10.1186/s12864-014-1206-5

